# Endosomal actin branching, fission and receptor recycling require FCHSD2 recruitment by MICAL-L1

**DOI:** 10.1101/2024.06.27.601011

**Authors:** Devin Frisby, Ajay B. Murakonda, Bazella Ashraf, Kanika Dhawan, Leonardo Almeida-Souza, Naava Naslavsky, Steve Caplan

## Abstract

Endosome fission is required for the release of carrier vesicles and the recycling of receptors to the plasma membrane. Early events in endosome budding and fission rely on actin branching to constrict the endosomal membrane, ultimately leading to nucleotide hydrolysis and enzymatic fission. However, our current understanding of this process is limited, particularly regarding the coordination between the early and late steps of endosomal fission. Here we have identified a novel interaction between the endosomal scaffolding protein, MICAL-L1, and the human homolog of the *Drosophila* Nervous Wreck (Nwk) protein, FCH and double SH3 domains protein 2 (FCHSD2). We demonstrate that MICAL-L1 recruits FCHSD2 to the endosomal membrane, where it is required for ARP2/3-mediated generation of branched actin, endosome fission and receptor recycling to the plasma membrane. Since MICAL-L1 first recruits FCHSD2 to the endosomal membrane, and is subsequently responsible for recruitment of the ATPase and fission protein EHD1 to endosomes, our findings support a model in which MICAL-L1 orchestrates endosomal fission by connecting between the early actin-driven and subsequent nucleotide hydrolysis steps of the process.

## Introduction

The trafficking of internalized cargos through endocytic compartments and their recycling to the plasma membrane (PM) is critical to cellular function and necessary for plasma membrane homeostasis and the regulation of cell signaling pathways (Naslavsky & Caplan, 2018). Indeed, higher-order processes that are regulated by membrane trafficking include membrane remodeling, signal transduction, cell migration, and control of cell polarity (Caswell & Norman, 2008; Cullen & Steinberg, 2018; Wang *et al*, 2000). Internalization at the plasma membrane occurs through various mechanisms of endocytosis, ultimately leading to the homotypic and heterotypic fusion of endocytic vesicles to generate the early or sorting endosome (EE/SE). At the EE/SE, cargos are actively sorted and shunted to different fates such as lysosomal degradation, retrograde trafficking to the Golgi, or recycling to the cell surface. Cargos destined for recycling are sorted and packaged into tubulovesicular structures that undergo fission and are either trafficked directly back to the plasma membrane (fast recycling) or first transported to the “endocytic recycling compartment,” a series of perinuclear tubular recycling endosomes, before moving to the plasma membrane (slow recycling) (Naslavsky & Caplan, 2018; Xie *et al*, 2016). While cargo recycling has previously been characterized as a passive or “default” process, recent studies support the notion that recycling is an active process mediated by cargo tail sorting signals and protein complexes that control recycling (Hsu *et al*, 2012; McNally & Cullen, 2018). For example, ACAP1 mediates the sorting and recycling of various receptors, including the transferrin receptor (TfR), Glut4, and β1-integrin (Bai *et al*, 2012; Dai *et al*, 2004; Li *et al*, 2005). Additionally, the retromer and retriever complexes cooperate with sorting nexins (SNXs) to regulate receptor sorting and recycling (Burd & Cullen, 2014; Cullen & Steinberg, 2018; McNally *et al*, 2017; Wang *et al*, 2018).

Both the retromer and retriever recycling complexes interact either directly or indirectly with the WASP and SCAR homolog (WASH) complex to activate ARP2/3-mediated branched actin polymerization (Gomez & Billadeau, 2009; Harbour *et al*, 2012; Jia *et al*, 2012; McNally *et al*., 2017). The WASH complex is recruited to endosomes via direct interaction with the retromer complex, while the retriever complex is indirectly recruited to endosomes by the WASH complex through an interaction with the COMMD/CCDC22/CCDC93 (CCC) complex (Gomez & Billadeau, 2009; Jia *et al*., 2012; McNally *et al*., 2017; Seaman *et al*, 2013). Interaction of the WASH complex with cargo recycling complexes highlights the role of actin in the early steps of cargo recycling at the endosome. Moreover, activation of ARP2/3-mediated branched actin polymerization establishes a physical barrier that helps form a cargo retrieval subdomain by inhibiting the diffusion of recycling receptors into the degradative endosomal subdomain (Puthenveedu *et al*, 2010; Simonetti & Cullen, 2019). In addition, treatment of cells with branched actin inhibitors decreased recycling endosome tubulation, further establishing a crucial role for actin in the tubulation of cargo-laden vesicles on endosomes (Anitei & Hoflack, 2011; Delevoye *et al*, 2016). During fission, actin provides a necessary pushing force at the neck of budding vesicles for constriction and generation of appropriate membrane tension (Derivery *et al*, 2009; Gomez & Billadeau, 2009). Through its regulation of receptor sorting, endosome tubulation, and endosome fission, the WASH complex is required for T cell receptor (TCR), GLUT1, β2AR, and a5β1-integrin recycling back to the plasma membrane; however, the proteins and mechanisms regulating actin polymerization and depolymerization at endosomes are largely unexplored (Piotrowski *et al*, 2013; Temkin *et al*, 2011; Zech *et al*, 2011).

Coordination of fission at endosomes is complex and incompletely understood. It requires the function of multiple proteins in addition to the WASH complex, ARP2/3, and branched actin. Several models have been proposed for fission at endosomes (Gopaldass *et al*, 2024; Naslavsky & Caplan, 2023a; Solinger & Spang, 2022). One recently elucidated mechanism for endosome fission is endoplasmic reticulum-driven fission, where branched actin patches containing Coronin1C stabilize cargo sequestration at tubular buds and recruit TMCC1 to drive fission (Hoyer *et al*, 2018; Rowland *et al*, 2014). However, it is unclear whether ER-based fission is primarily a mechanism for endosome homeostasis or whether it is key to vesicle/tubule carrier release and transport for recycling. Amphipathic helix insertion and the induction of positive membrane curvature is another mechanism that has been recently identified for endosome fission (Courtellemont *et al*, 2022; Gopaldass *et al*, 2017). Many models, however, favor a combination of initial actin-based membrane constriction followed by nucleotide hydrolysis of a dynamin-family fission protein to detach the budding vesicle/tubule ((Derivery *et al*., 2009) and reviewed in (Naslavsky & Caplan, 2018, 2023a)).

The mode by which the earlier actin-based steps of endosomal membrane constriction are linked to the terminal steps of fission remain poorly defined. One crucial endosomal membrane hub and scaffolding protein is MICAL-L1 (Fig. 1A). MICAL-L1 appears to be a master regulator of endosome fission; it localizes to endosomes and recruits a variety of proteins involved in both the early and late steps of endosome fission, including Syndapin2/PACSIN2 and EHD1 (Giridharan *et al*, 2013; Sharma *et al*, 2009). Syndapins may link membrane trafficking to cortical actin cytoskeletal dynamics, as they interact with dynamin and N-WASP (Qualmann *et al*, 1999). EHD1 has been implicated in the later stages of fission (Cai *et al*, 2012; Cai *et al*, 2013; Cai *et al*, 2014; Deo *et al*, 2018; Dhawan *et al*, 2020; Kamerkar *et al*, 2019; Naslavsky & Caplan, 2011). Indeed, depletion of either MICAL-L1 or EHD1 leads to impaired fission and recycling (Cai *et al*., 2012; Cai *et al*., 2013; Cai *et al*., 2014; Deo *et al*., 2018; Dhawan *et al*., 2020; Farmer *et al*, 2020; Kamerkar *et al*., 2019; Naslavsky & Caplan, 2011), and evidence suggests that EHD1 ATP hydrolysis drives its oligomerization on the endosomal membrane to induce membrane thinning and fission (Deo *et al*., 2018; Sharma *et al*., 2009). How MICAL-L1 coordinates the actin polymerization and membrane tubulation at endosomes that occurs during the early stages of cargo sorting with the later stages of EHD1-mediated endosome fission remains unknown.

**Figure 1.**
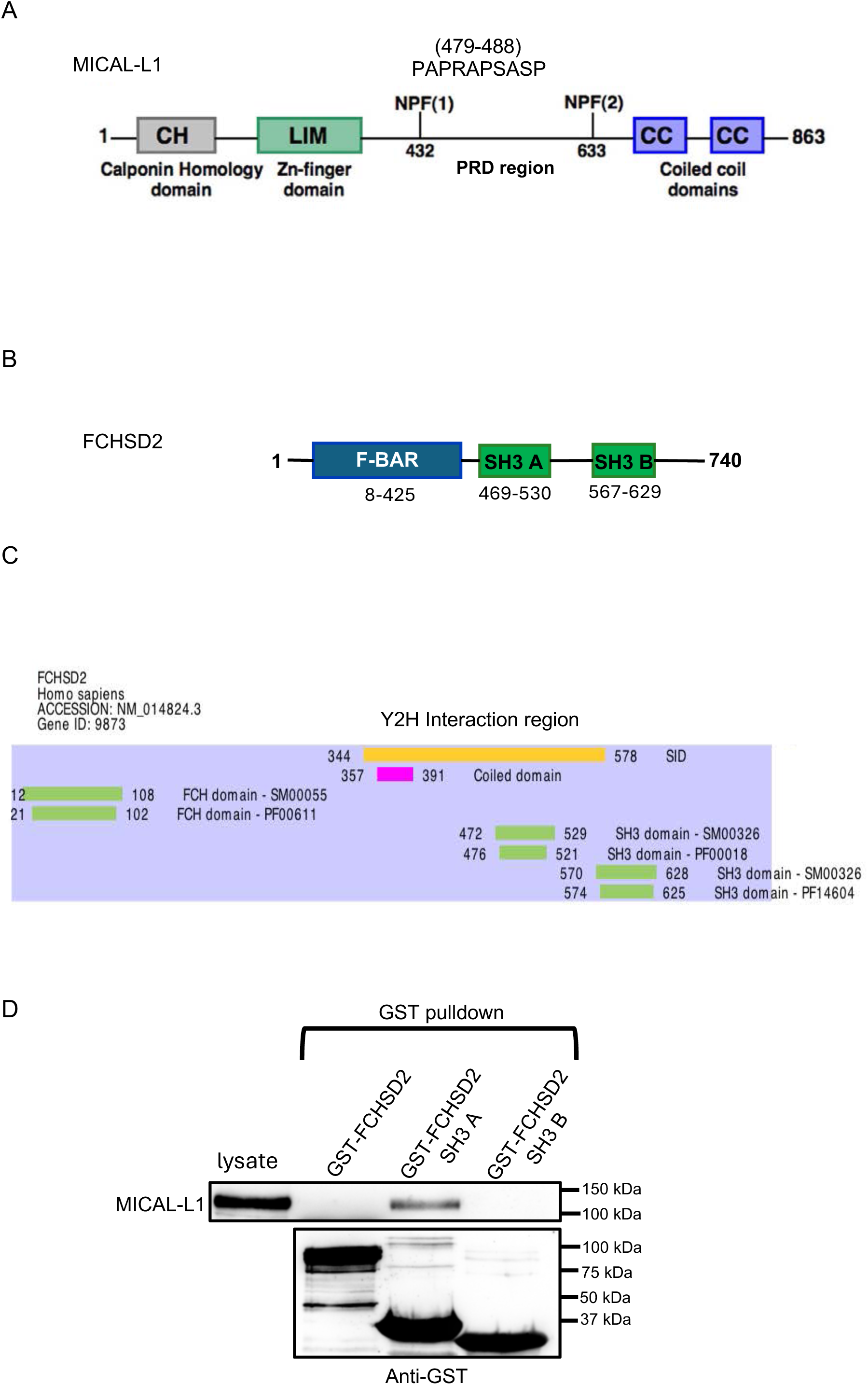
FCHSD2 interacts with MICAL-L1 via its first SH3 domain. A. Domain architecture of MICAL-L1. B. Domain architecture of FCHSD2. C. An unbiased yeast 2-hybrid screen identified FCHSD2 as a MICAL-L1 interactor. MICAL-L1 was used as bait and an area covered by FCHSD2 amino acids 344-578 (yellow region, encompassing the first SH3 domain) was identified in the interaction. D. GST-pulldowns identified FCHSD2 SH3 A as the MICAL-L1 interacting domain. GST-FCHSD2, GST-FCHSD2 SH3 A, and GST-FCHSD2 SH3 B proteins were expressed in bacteria and purified. Purified protein (20 μg) was incubated with HeLa lysate and pulled down with Glutathione Sepharose resin. Samples were eluted, subjected to SDS-PAGE and immunoblotted with anti-GST and anti-MICAL-L1 primary antibodies. MICAL-L1=Microtubule Associated Monooxygenase, Calponin and Lim Domain Containing, FCHSD2=FCH and Double SH3 Domains 2, CH=Calponin Homology, LIM=Lin11, Islet-1 and Mec-3, CC=coiled coil, BAR=Bin-Amphiphysin-Rvs, SH3=Src Homology 3, Y2H=Yeast 2-Hybrid, GST=Glutathione S-Transferase.

To address this question, we took advantage of an unbiased yeast 2-hybrid screen and identified FCH and double SH3 domains protein 2 (FCHSD2) as a novel MICAL-L1 interacting partner. FCHSD2 contains an N-terminal F-BAR domain, two SRC homology 3 (SH3) domains, and a C-terminal proline-rich region (Fig. 1B). Initial insights into the function of FCHSD2 come from studies done in fruit flies with the *Drosophila* homolog, Nervous Wreck (Nwk). Adult temperature-sensitive Nwk mutants become paralyzed and experience spasms at 38°C (Coyle *et al*, 2004). Investigations into the function of Nwk reveal that it regulates receptor signaling at neuromuscular junctions (NMJs) through its cooperation with the endocytic proteins Dap160 and WASP and the regulation of Cdc42/WASP-mediated branched actin polymerization (O’Connor-Giles *et al*, 2008; Rodal *et al*, 2008). FCHSD2 interacts with the corresponding mammalian homologs of Dap160 and WASP, intersectin-1 (ITSN-1) and N-WASP (Almeida-Souza *et al*, 2018), respectively, as well as with dynamin (O’Connor-Giles *et al*., 2008; Rodal *et al*., 2008) and SNX9/18 (Haberg *et al*, 2008), and FCHSD2 knockdown impedes cargo internalization via clathrin-mediated endocytosis (CME) (Almeida-Souza *et al*., 2018; Xiao *et al*, 2018). Indeed, FCHSD2 regulates CME by enhancing N-WASP/ARP2/3-mediated branched actin polymerization to promote maturation of clathrin-coated pits and facilitate dynamin-mediated fission and internalization (Almeida-Souza *et al*., 2018). More recently, Nwk and FCHSD2 have been implicated in the regulation of cargo recycling at endosomes. Nwk localizes to Rab11-containing endosomes in *Drosophila* NMJs and regulates receptor trafficking through direct interaction with sorting nexin 16 (Rodal *et al*, 2011; Rodal *et al*., 2008), and Nwk mutants phenocopy RAB11 mutants for extracellular vesicle cargo transport (Blanchette *et al*, 2022). Interestingly, FCHSD2 knockdown interferes with transferrin (Tf) and epidermal growth factor receptor (EGFR) recycling and subsequently leads to increased localization of these cargos with LAMP1, a marker of the lysosomal membrane (Xiao & Schmid, 2020). However, the mechanism by which FCHSD2 regulates cargo recycling at endosomes remains unknown.

Here we identify FCHSD2 as a novel MICAL-L1 interacting partner. MICAL-L1 knock-out impairs FCHSD2 localization to endosomes. Consistent with previous findings, FCHSD2 depletion had a minor but significant decrease on uptake of both clathrin-dependent and - independent cargo. Importantly, FCHSD2 knock-down impaired the recycling of cargos from both internalization pathways. We show that upon FCHSD2 depletion, the size of EEA1- and MICAL-L1-decorated endosomes increases, and we demonstrate that FCHSD2 knockdown significantly impedes endosomal fission. Moreover, FCHSD2 depletion decreases the concentration of branched actin at endosomes, supporting its function in activating branched actin polymerization and fission at endosomes.

## Results

### MICAL-L1 interacts with and recruits FCHSD2 to endosomes

The execution of fission at the endosome is a complex process that requires tightly coordinated orchestration of multiple proteins in a sequential manner. MICAL-L1 (Fig. 1A) is a key scaffold that, as described above, interacts with multiple endosomal proteins. However, a central question is how this scaffold coordinates the fission process and how it links nucleotide hydrolysis-driven fission with the earlier steps of membrane budding and constriction.

To identify novel MICAL-L1 interaction partners that might play a role in endosome fission, we used an unbiased yeast two-hybrid approach to screen over 100,000,000 potential interactions with full-length MICAL-L1 as bait (Fig. 1A). Among the potential hits identified with moderate confidence was the human homolog of the *D. melanogaster* Nervous Wreck (Nwk) protein (Becalska *et al*, 2013; Coyle *et al*., 2004; Rodal *et al*., 2011; Rodal *et al*., 2008), known as FCH and double SH3 domains 2 (FCHSD2; also known as CAROM) (Almeida-Souza *et al*., 2018; Ohno *et al*, 2003; Xu *et al*, 2017) (Fig. 1B). Although primarily known for its role in internalization at the plasma membrane (Almeida-Souza *et al*., 2018; Xiao & Schmid, 2020), recent studies have implicated mammalian FCHSD2 in receptor recycling (Xiao *et al*., 2018; Xiao & Schmid, 2020). FCHSD2 has an F-BAR domain as well as two SH3 domains. Indeed, the binding region identified through the yeast two-hybrid screen included the first of the two FCHSD2 SH3 domains, but not the second SH3 domain (Fig. 1C). To validate this interaction, we generated and purified GST fusion proteins for full-length human FCHSD2 (GST-FCHSD2) and GST fused to either the first or second SH3 domain (GST-SH3 A and GST-SH3 B). Upon incubation with HeLa cell lysates, we demonstrated that whereas GST-FCHSD2 and GST-SH3 B were unable to pull-down MICAL-L1, the isolated GST-SH3 A domain precipitated MICAL-L1 (Fig. 1D). These data are consistent with a role for SH3 A in binding to MICAL-L1, likely via one or more of the latter’s proline-rich motifs. The lack of binding for GST-FCHSD2 supports the notion that the full-length protein is in an autoinhibited conformation (Almeida-Souza *et al*., 2018; Del Signore *et al*, 2021; Rodal *et al*., 2008; Stanishneva-Konovalova *et al*, 2016).

The localization of FCHSD2 in mammalian cells has been studied primarily using exogenous protein, with reports indicating subcellular localization underneath the plasma membrane (Almeida-Souza *et al*., 2018) and at actin protrusions (Zhai *et al*, 2022b). Given the role of MICAL-L1 in scaffolding endosome fission machinery, we hypothesized that some FCHSD2 might be transiently localized to endosomes. To evaluate if low concentrations of FCHSD2 are detectable on endosomes, we overexpressed GFP-FCHSD2 together with the active GTP-locked RAB5 mutant Q79L (Stenmark *et al*, 1994). As expected, the RAB5 mutant induced and localized to enlarged endosomal structures (Fig. 2A; inset in Fig. 2B). Strikingly, FCHSD2 could be detected at the surface of these enlarged endosomes (Fig. 2C; inset in 2D, and merged in Fig. 2E and F). To quantify the overlap between FCHSD2 and RAB5, we performed line scans on multiple images. We measured the relative intensity of the fluorescence for each channel along the line, normalizing the fluorescence by subtracting the “background” fluorescence from the cytoplasm. As demonstrated, the graph shows a peak of mean fluorescence for each channel, with the peak FCHSD2 (green) intensity overlapping with the peak RAB5 (red) intensity (Fig. 2G). These data support the notion that FCHSD2 can be detected in proximity to RAB5 on the endosomal membrane.

**Figure 2.**
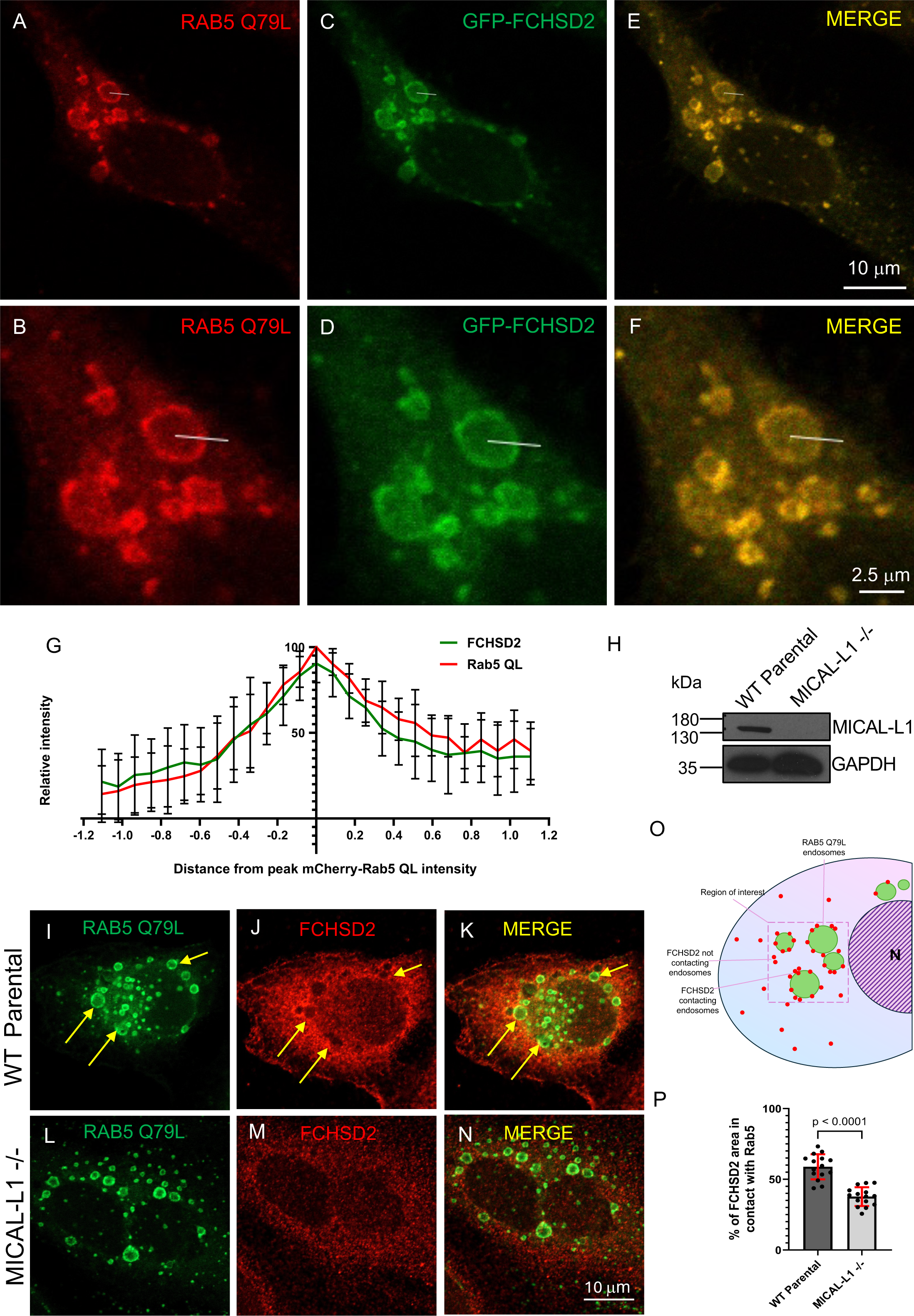
FCHSD2 is recruited to Rab5 QL endosomes by MICAL-L1. A-F. HeLa cells were co-transfected with GFP-FCHSD2 and the mCherry-RAB5 Q79L mutant, which remains GTP-locked and active. Cells were fixed, imaged, and analyzed in ImageJ. B, D and F represent insets of A, C and E, respectively. A profile 2.81 µm in length was drawn from the cytoplasm into the enlarged mCherry-RAB5 QL vesicles, and the fluorescence intensities of both channels was collected along the line. Background subtraction was performed and the fluorescence intensities along each profile were normalized. G. The data (quantified from from A-F) were plotted as relative intensity over the distance from peak mCherry-Rab5 QL intensity (red) and demonstrates that FCHSD2 staining overlaps with RAB5 QL on the endosomal membrane. H. Immunoblot validation of MICAL-L1 knock-out in the CRISPR/Cas9 gene-edited knock-out cell line. I-N. HeLa WT parental or MICAL-L1 knockout cells were transfected with the GFP-RAB5 Q79L mutant. Cells on coverslips were fixed after transfection and immunostained with anti-FCHSD2 to visualize endogenous FCHSD2 (J,M). Confocal images were captured and analyzed in Imaris software. As demonstrated, endogenous FCHSD2 coats RAB5 Q79L endosomes in parental cells (I-K; yellow arrows) but is largely absent from the endosomes in MICAL-L1 knockout cells (L-N). O. Segmentation strategy for quantification of the percentage of cortactin in contact with RAB5 Q79L endosomes. Square regions of interest (ROI) were made to include the maximal endosomal area within the ROI. P. Quantification of I-N. In a demarked ROI around the RAB5 Q79L endosomes, the volume of FCHSD2 puncta that made contact with GFP-RAB5 Q79L endosomes was represented as a percentage of the total FCHSD2 volume in that region.

Given that MICAL-L1 binds directly with lipids such as phosphatidic acid and phosphatidylserine (Giridharan *et al*., 2013) and interacts with FCHSD2, we next hypothesized that MICAL-L1 is responsible for the initial recruitment of FCHSD2 to endosomes. To address this, we again used the RAB5 Q79L mutant to transfect either parental or MICAL-L1^-/-^ HeLa cells (CRISPR/Cas9 knock-out validated in Fig. 2H). We first confirmed that endogenous FCHSD2 could be detected coating the membrane of enlarged RAB5 Q79L endosomes in the parental cells (Fig. 2I-K; quantified in P). However, using an unbiased semi-automated quantification method (see Fig. 2O for outline of segmentation method), significantly less FCHSD2 was observed on the RAB5 Q79L endosomes in the MICAL-L1^-/-^ cells (Fig. 2L-N; quantified in P). These data support the idea that MICAL-L1 is required for recruitment of FCHSD2 to the endosomal membrane.

### FCHSD2 is required for internalization and recycling of clathrin-dependent and -independent cargo

Since MICAL-L1 has been implicated as an endosomal regulator that is required for receptor recycling and FCHSD2 can be detected on endosomes, we next tested whether FCHSD2 depletion impairs recycling of cargo from endosomes. We first assessed the impact of FCHSD2 depletion on transferrin receptor (TfR) which is internalized through clathrin-mediated endocytosis. After 10 min. of transferrin uptake, FCHSD2-depleted cells (validated in Fig. 3E) displayed a modest decrease in internalized TfR of about 10%, consistent with previous studies (Almeida-Souza *et al*., 2018; Xiao & Schmid, 2020) (Fig. 3A and C; quantified in Fig. 3F). However, after a chase of 50 min. to follow TfR recycling and the exit of labeled transferrin from the cell, the FCHSD2 knock-down cells displayed a significant delay in endocytic recycling (normalized to levels of internalized TfR) (Fig. 3B and D; quantified in Fig. 3G). These delays in recycling were consistent with those reported by Xiao and Schmid for both TfR and epidermal growth factor receptor (Xiao & Schmid, 2020).

**Figure 3.**
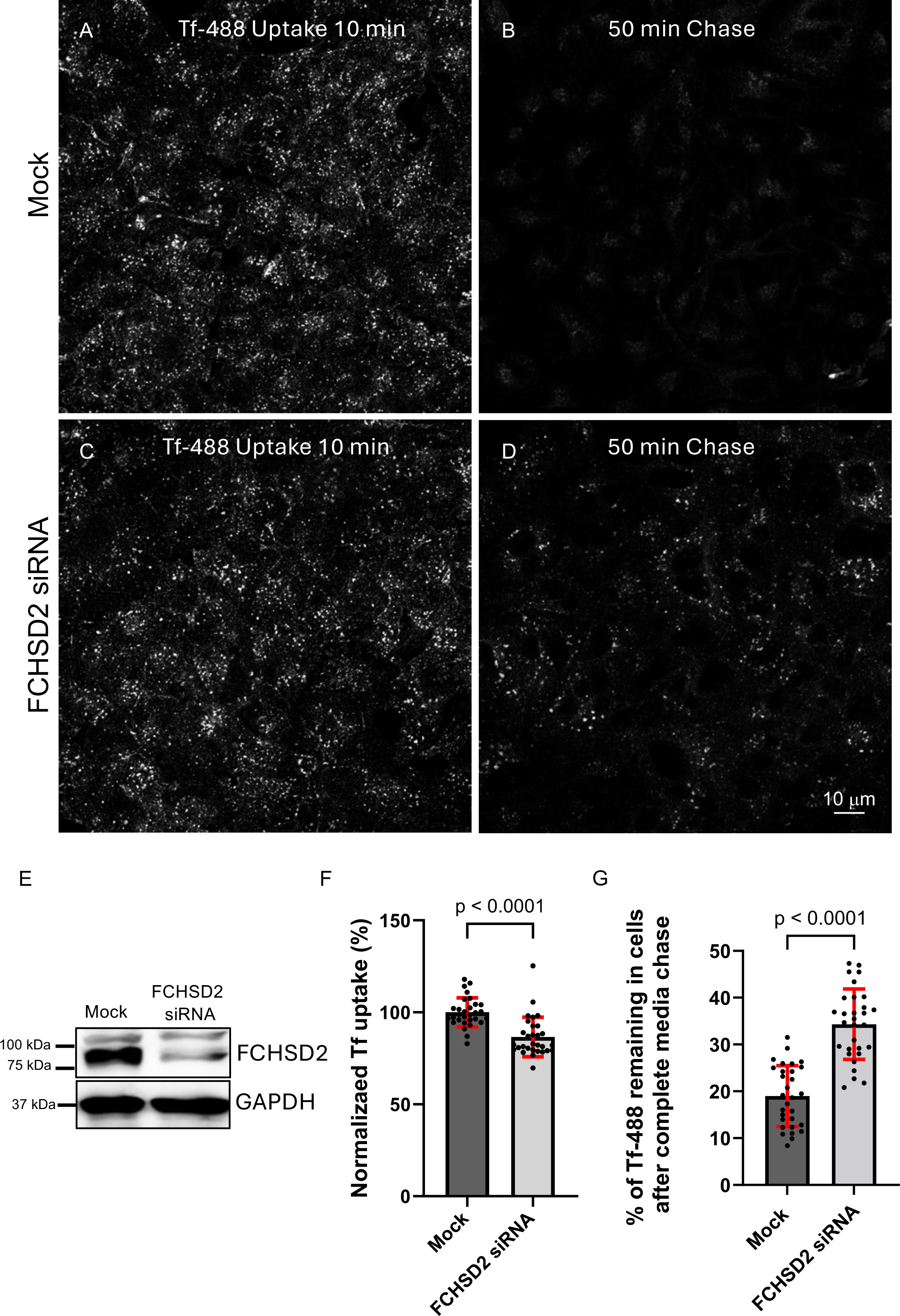
Transferrin uptake and recycling are impaired upon FHCSD2 depletion. A-D. Mock and FCHSD2 siRNA knock-down HeLa cells were incubated with fluorophore-labeled transferrin (Tf-488) for 10 min (A,C) and chased with complete media for 50 min to allow recycling (B,D). E. Immunoblot validation of FCHSD2 siRNA knock-down. F. FCHSD2 depletion impairs transferrin uptake. Images (similar to those in A and C) were analyzed in Zeiss Zen Blue software by measuring the arithmetic mean intensity of each image after uptake, which was plotted relative to the highest arithmetic mean intensity. G. FCHSD2 depletion delays transferrin recycling. Images (similar to those in B and D) were analyzed in Zeiss Zen Blue software by measuring the arithmetic mean intensity of each image after recycling. These values were normalized to the mean arithmetic mean intensity after uptake.

To determine if FCHSD2 is involved in the recycling of receptors internalized through clathrin-independent means, we next analyzed the internalization and recycling of major histocompatibility complex class I (MHC I) receptors (Fig. 4). Using both FCHSD2 siRNA knock-down and FCHSD2^-/-^ cells (validated in Fig. 4K), we demonstrated that whereas FCHSD2 depleted cells showed a modest decrease in internalized MHC I (Fig. 4A-D; quantified in Fig. 4I), both FCHSD2 knock-down and knock-out cells displayed delays in MHC I recycling, with the “acute” siRNA knock-down cells displaying more severe recycling defects than the “chronic” FCHSD2^-/-^ cells (Fig. 4E-H; quantified in Fig. 4J). Overall, our data support the notion that FCHSD2 plays an important role in regulating events at the endosome that are required for receptor recycling.

**Figure 4.**
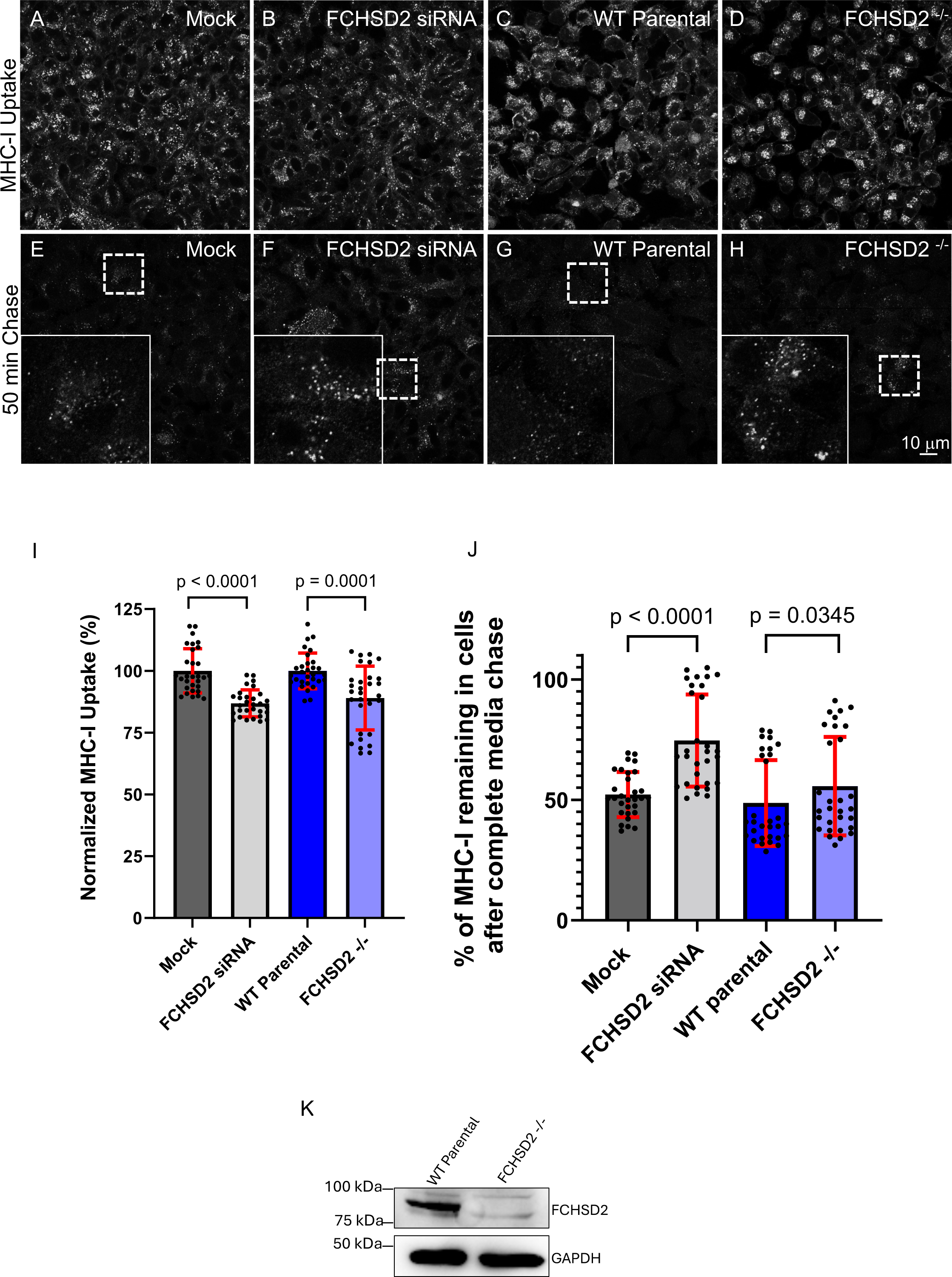
MHC-I uptake and recycling are impaired upon FCHSD2 depletion. A-D. Mock, FCHSD2 siRNA knock-down, WT parental, and FCHSD2 knock-out HeLa cells were incubated with anti-MHC1 antibodies for 20 min (A-D). Cells were glycine stripped to remove non-internalized antibody, fixed, immunostained, and imaged. E-H. Following the 20 min uptake, cells were glycine stripped, washed, and chased with complete DMEM for 50 min to allow for MHC-I recycling. After chase, cells were glycine stripped again, fixed, and immunostained to detect non-recycled MHC-I. Demarked regions are represented as insets. I. FCHSD2 depletion impairs MHC-I uptake. Images (including A-D) were analyzed in Zeiss Zen Blue software by measuring the arithmetic mean intensity of each image after uptake, which was plotted relative to the highest arithmetic mean intensity. J. FCHSD2 depletion impairs MHC-I recycling. Images (including E-H) were analyzed in Zeiss Zen Blue software by measuring the arithmetic mean intensity of each image after recycling. These values were normalized to the mean arithmetic mean intensity after uptake. K. Immunoblot validation of FCHSD2 CRISPR/Cas9 knock-out in the CRISPR/Cas9 gene-edited knock-out cell line (FCHSD2^-/-^).

There is a growing appreciation that many endocytic regulatory proteins are also involved in regulating primary ciliogenesis (Bales & Gross, 2016; Madhivanan & Aguilar, 2014; Pedersen *et al*, 2016), including MICAL-L1 (Xie *et al*, 2019; Xie *et al*, 2023). Accordingly, we asked whether FCHSD2 depletion impacts the ability of retinal pigmented epithelial cells (RPE-1) to generate a primary cilium. Upon FCHSD2 knock-down (validated in Fig. EV1 C), a greater percentage of serum-starved RPE-1 cells generated a primary cilium compared to mock-treated cells (Fig. EV1, compared B to A; quantified in D). These data are consistent with a role for FCHSD2 in endocytic regulatory function and actin regulation, which has been linked to inhibition of ciliogenesis (Hoffman & Prekeris, 2022).

### FCHSD2 is involved in the endosomal fission process

The fission of endosomes is required for formation of carrier vesicles/tubules and the recycling of receptors to the plasma membrane. Given the ascribed role for MICAL-L1 in endosome fission (Cai *et al*., 2014; Farmer *et al*., 2020; Rahajeng *et al*, 2012; Sharma *et al*., 2009), we postulated that FCHSD2 may regulate fission at endosomes, potentially through its actin-regulatory activity. In recent years, it has become evident that in addition to punctate endosomes and endosomal carriers, many endosomes and endosomal carriers are tubular and their fission is regulated by MICAL-L1 and its interaction partners (Dhawan *et al*., 2020; Dhawan *et al*, 2022; Farmer *et al*, 2021; Jones *et al*, 2020). Accordingly, we asked whether the fission of punctate and tubular-shaped endosomes is impacted by depletion of FCHSD2 (Fig. 5 and EV2). As shown, depletion of FCHSD2 by siRNA knock-down and in FCHSD2^-/-^ cells (Fig. 5B and D) led to enlarged EEA1-marked endosomes (compare Fig. 5B to A and D to C; quantified in E). FCHSD2 knock-down also led to a longer and more elaborate tubular endosome network as compared to mock-treated cells (EV2, compare B to A; quantified in EV2 H). Similarly, FCHSD2^-/-^ cells also displayed an increase in total area of tubular endosomes marked by MICAL-L1 compared to the wild-type parental cells (EV2 compare D to C; quantified in EV2 I). In addition, we transfected the FCHSD2^-/-^ cells with wild-type FCHSD2. As shown, the FCHSD2-transfected cells (marked by red stars) displayed fewer tubular endosomes than untransfected cells (tubular MICAL-L1 endosomes marked by green arrows) (EV2 E-G), and quantification showed a similar mean tubular endosome area to that observed in the wild-type parental cells (quantified in EV2 I). These data support the idea that FCHSD2 functions in the fission of EEA1 punctate endosomes and MICAL-L1-marked tubular endosomes.

**Figure 5:**
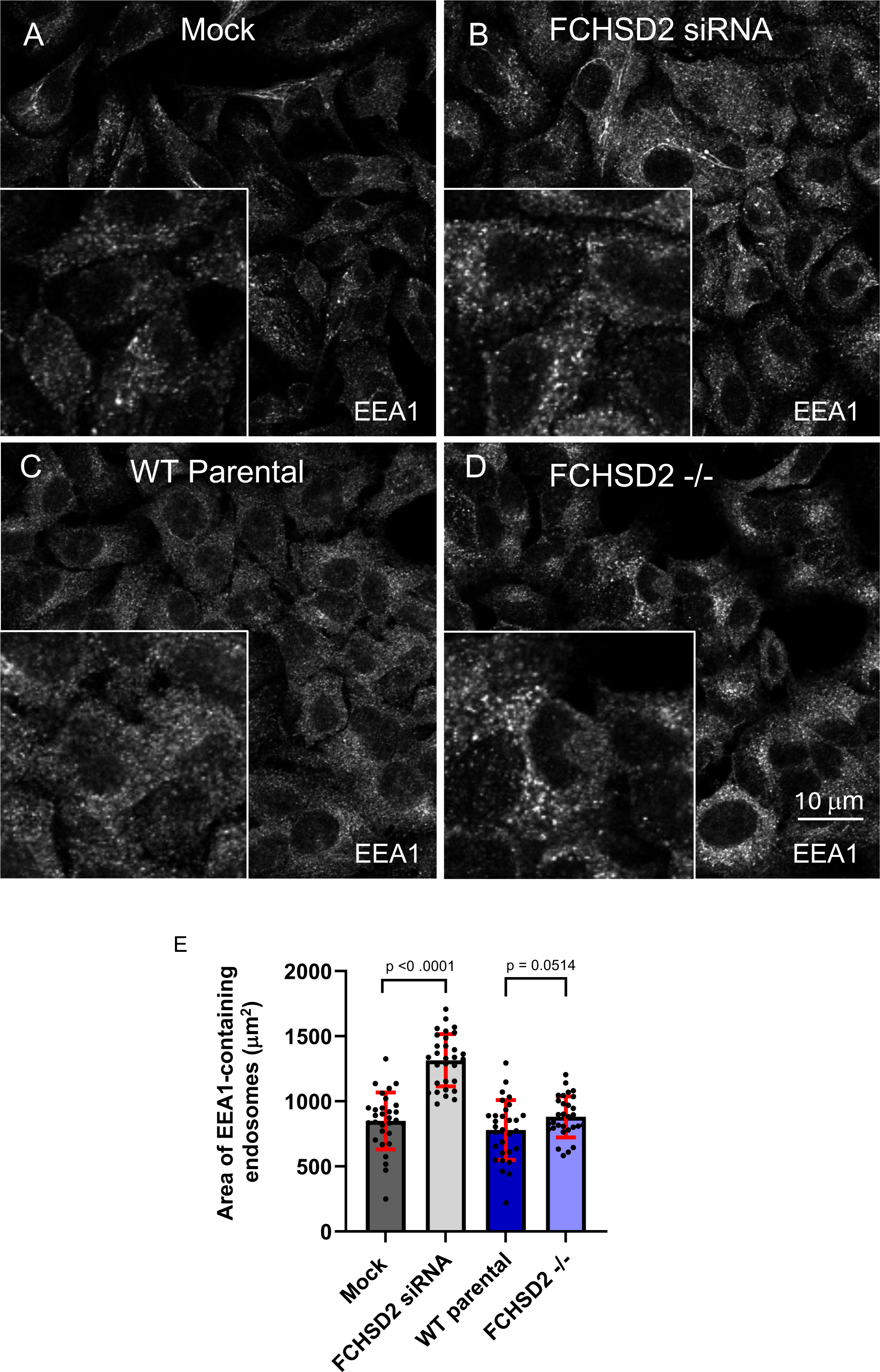
FCHSD2 depletion leads to increased endosome size. A-D. Mock-treated (A), FCHSD2 knock-down (B), WT parental (C), and FCHSD2^-/-^(D) cells were fixed and immunostained with a primary antibody against the endosome marker EEA1. Insets depict the increased size of EEA1-marked endosomes. E. Quantification of the increased EEA1 area per cell shown in A-D. Using Imaris software, EEA1 endosomes were rendered as surfaces, and the EEA1 area per cell was calculated per image and plotted. Calculations are derived from 30 images from 3 independent experiments.

While endosome size usually correlates negatively with fission in cells, size is also affected by fusion. To more definitively quantify the effects of FCHSD2 depletion on endosome fission, we took advantage of a novel in-cell fission assay that we recently developed that uses a synchronized system to acutely measure decrease in endosome size over a 30 min. period (Dhawan *et al*., 2022). Briefly, in mock and FCHSD2 knock-down cells, internalized transferrin was used to mark endosomes, and both mock and knock-down cells were incubated with the PI3K inhibitor LY294002 to induce enlarged endosomes and allow synchronization of fission events upon inhibitor washout (chase). After inhibitor washout, in both mock-treated and FCHSD2-depleted cells, we imaged transferrin-containing endosomes in 3D and quantified the mean size of more than 100,000 structures, measuring the frequency of structures from the binned endosome sizes (Fig. 6A-D; quantified in Fig. 6E-G). By integrating the area below each curve and subtracting the values from before and after the LY294002 washout (chase), we calculated and plotted a mean value for fission (Fig. 6G). Additional experiments were done to measure the size of EEA1 endosomes (rather than transferrin-containing endosomes), and they also demonstrated whereas mock-treated cells displayed reduced endosome size after LY294002 washout, FCHSD2 knock-down cells showed no significant reduction in size, suggesting impaired endosome fission (EV3). Overall, these data strongly support a role for FCHSD2 in endosome fission.

**Figure 6:**
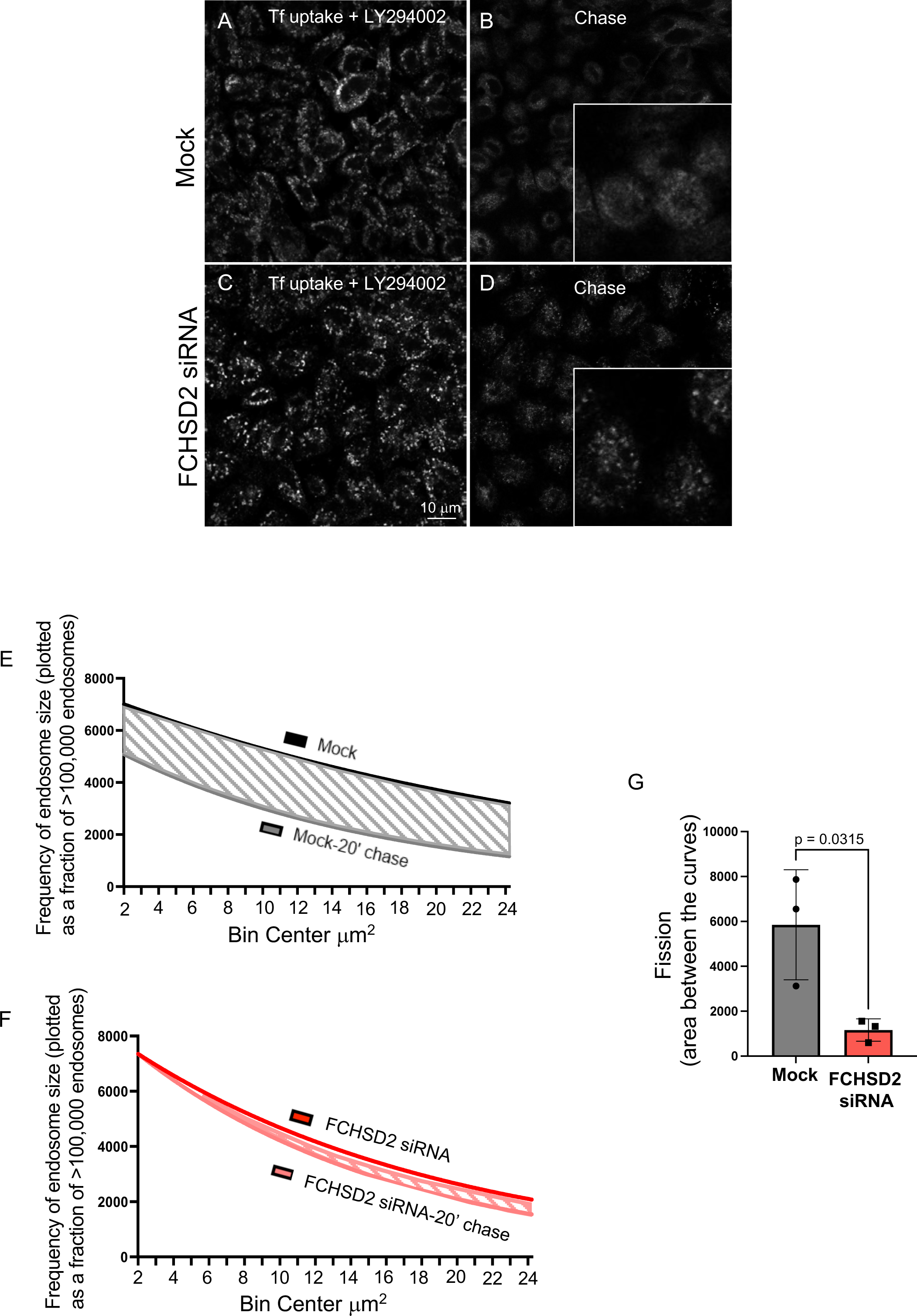
FCHSD2 depletion leads to impaired endosome fission. A-D. Cells on coverslips were treated with the PI3K inhibitor LY294002 for 45 min to induce enlarged endosomes and synchronize the size of the endosome population. The cells were then incubated with Tf-488 in the presence of the inhibitor for 15 additional minutes. Cells were immediately fixed (uptake; A,C) or chased with complete media to washout the inhibitor and allow fission and recycling for 20 min (chase; B,D). E-F. Imaris software was used to quantify and bin hundreds of thousands of endosomes according to mean size. The surface function in Imaris was used to render all of the Tf-containing structures. The surface areas of over 100,000 Tf-containing vesicles were plotted as a frequency distribution plot in GraphPad Prism. A frequency distribution (interleaved) graph with bins from 2 μm up to 10 μm and the bin width set at 1 μm was plotted, and a Gaussian curve was extrapolated beyond 10 μm to infinity on the frequency distribution graphs for all four experimental groups. I. G. The area between the curves was calculated in GraphPad Prism by taking the difference between the area underneath the curves for uptake and chase in both experimental groups (E and F). The area between the curves represents a difference in the size of the endosomes after complete media chase, suggesting decreased fission in the FCHSD2 siRNA knockdown cells.

### FCHSD2 generates branched actin at the endosomal membrane

We next addressed the potential mechanism by which FCHSD2 regulates endosome fission. Given the role of FCHSD2 in actin regulation and control of WASP (Almeida-Souza *et al*., 2018; Becalska *et al*., 2013; Rodal *et al*., 2008; Stanishneva-Konovalova *et al*., 2016; Zhai *et al*., 2022b), we hypothesized that FCHSD2 may regulate fission by activation of ARP2/3 leading to actin polymerization and branching at the endosomal membrane, thus facilitating membrane budding. Indeed, treatment of cells with the ARP2/3 inhibitor, CK-666 (Nolen *et al*, 2009), led to impaired actin filament generation at RAB5 Q79L enlarged endosomes, whereas the CK-689 control had no effect on endosomal actin (EV4), highlighting the role of ARP2/3 in generating branched actin at endosomes.

To determine if FCHSD2 is required for ARP2/3-mediated endosomal branched actin, we used a modification of the elegant assay originally described by Cooper and colleagues (Zhao *et al*, 2013) and applied on endosomes (Muriel *et al*, 2016). Endogenous EEA1 was used as a marker of endosomes, and cortactin was used to identify branched actin in mock and FCHSD2 knock-down cells (Fig. 7). We then segmented cells into “peripheral” and “perinuclear” regions. As demonstrated for the peripheral segmentations and quantification, mock-treated cells displayed a significantly higher percent of endosomes with branched actin marked by cortactin (Fig. 7C-E; see arrows in E-inset) than FCHSD2 knock-down cells (Fig. 7F-H and insets; quantified in Fig. 7A). Contact between endosomes and cortactin in the perinuclear region (Fig. 7B) is also modestly decreased, but given the high density of endosomes in that region, the periphery provides a better opportunity for quantification.

**Figure 7.**
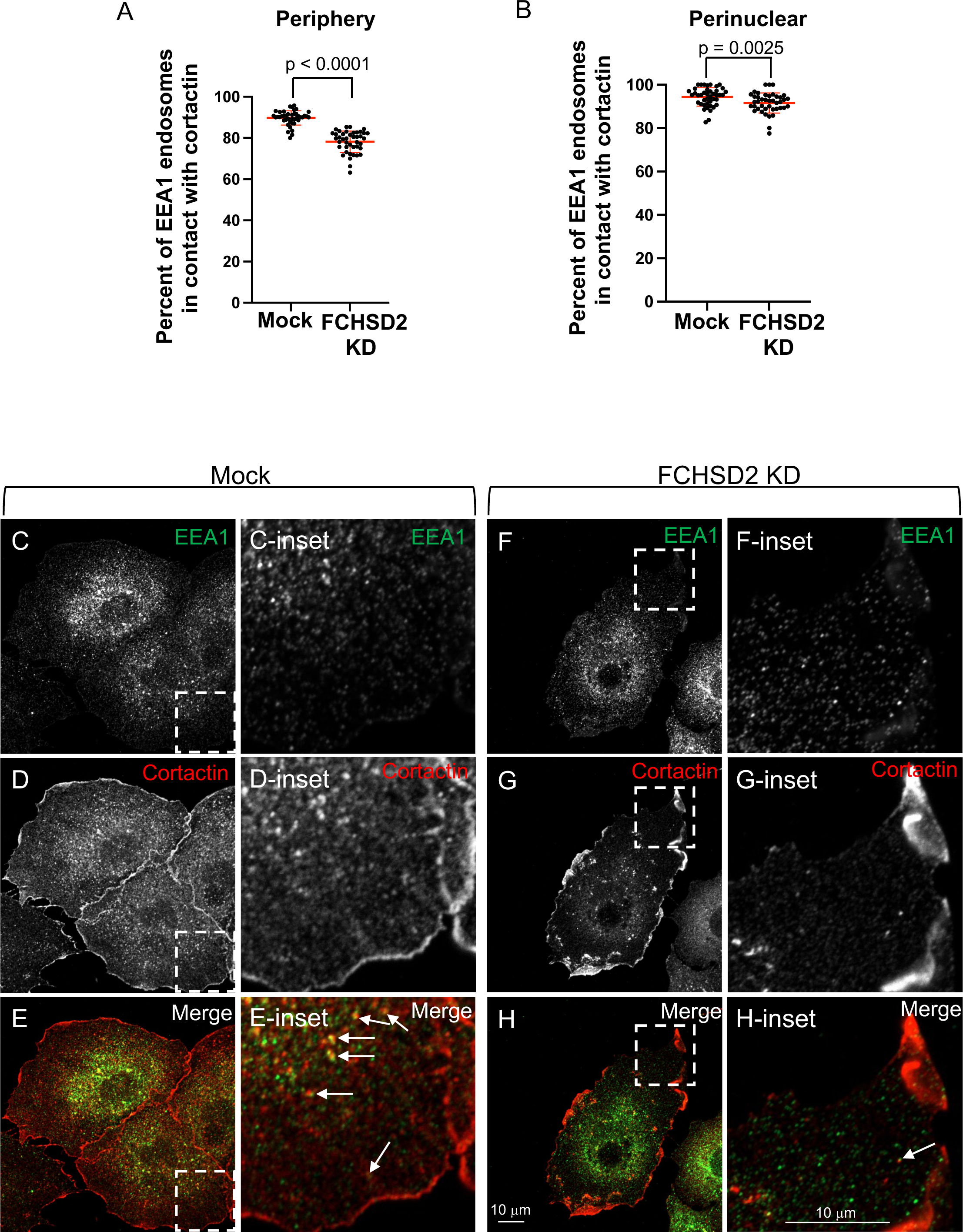
FCHSD2 knock-down leads to decreased branched actin at endosomes. A,B. Quantification of C-H. Three regions of interest (ROI) were demarked in the periphery of the cells (ROI begins at a minimal distance of 5 μm from the nucleus), and another 3 regions in the perinuclear area (ROI begins within 1 μm from the nucleus). The surfaces function in Imaris was used to 3D render all of the EEA1 and cortactin structures. EEA1 structures contacting cortactin were defined as EEA1 surfaces with a shortest distance to a cortactin surface with a value equal to zero. The EEA1 structures that contacted cortactin were represented as a percentage of the total number of EEA1 structures in either the perinuclear region or the cell periphery. C-E. Mock-treated NSCLC cells were fixed and immunostained with antibodies against cortactin (red) and EEA1 (green). Confocal images were captured and quantified (A,B). Insets of peripheral regions are shown. F-H. FCHSD2 knock-down cells were fixed and immunostained with antibodies against cortactin (red) and EEA1 (green). Confocal images were captured and quantified (A,B). Insets of peripheral regions are shown. EEA1-decorated endosomes in mock-treated NSCLC cells contact cortactin puncta (white arrows) more frequently than the FCHSD2 knockdown cells.

To better visualize actin and branched actin at endosomes in the presence and absence of FCHSD2, we transfected parental and FCHSD2 knock-out cells with the GTP-locked RAB5 Q79L mutant, and then immunostained for branched actin (cortactin) (Fig. 8). In parental cells, RAB5 Q79L induced formation of enlarged endosomes that were positive for cortactin and branched actin, often observed in a polarized manner at one or more region of the endosomal membrane (Fig. 8A-C, see arrows; quantified in Fig. 8P). Strikingly, in FCHSD2 knock-out cells branched actin and actin filaments were largely absent from the enlarged endosomes (Fig. 8D-F; quantified in Fig. 8P). However, when FCHSD2^-/-^ cells were rescued by transfection of WT FCHSD2, the levels of cortactin observed on the enlarged endosomes increased and were more similar to those in the parental cells than the FCHSD2^-/-^ cells (Fig. 8G-I; quantified in Fig. 8P). Transfection with a FCHSD2 Y478A/Y480A SH3 A mutant, that has impaired binding to proline rich motifs (Almeida-Souza *et al*., 2018) largely failed to rescue cortactin localization to endosomes (Fig. 8J-L; quantified in Fig. 8P). However, transfection with FCHSD2 Y576S+F607S, an SH3 B interface mutant that fails to interact with intersectin 1 (Almeida-Souza *et al*., 2018), nonetheless displayed significant rescue of branched actin generation at endosomes, similar to that observed upon rescue with the WT FCHSD2 (Fig. 8M-O; quantified in Fig. 8P). MICAL-L1^-/-^ cells were similarly devoid of branched actin (EV5). These data support the notion that MICAL-L1 recruits FCHSD2 by binding to its SH3 A domain to play a role in receptor recycling by activation of ARP2/3, a process which is required for the budding and fission of endosomes.

**Figure 8.**
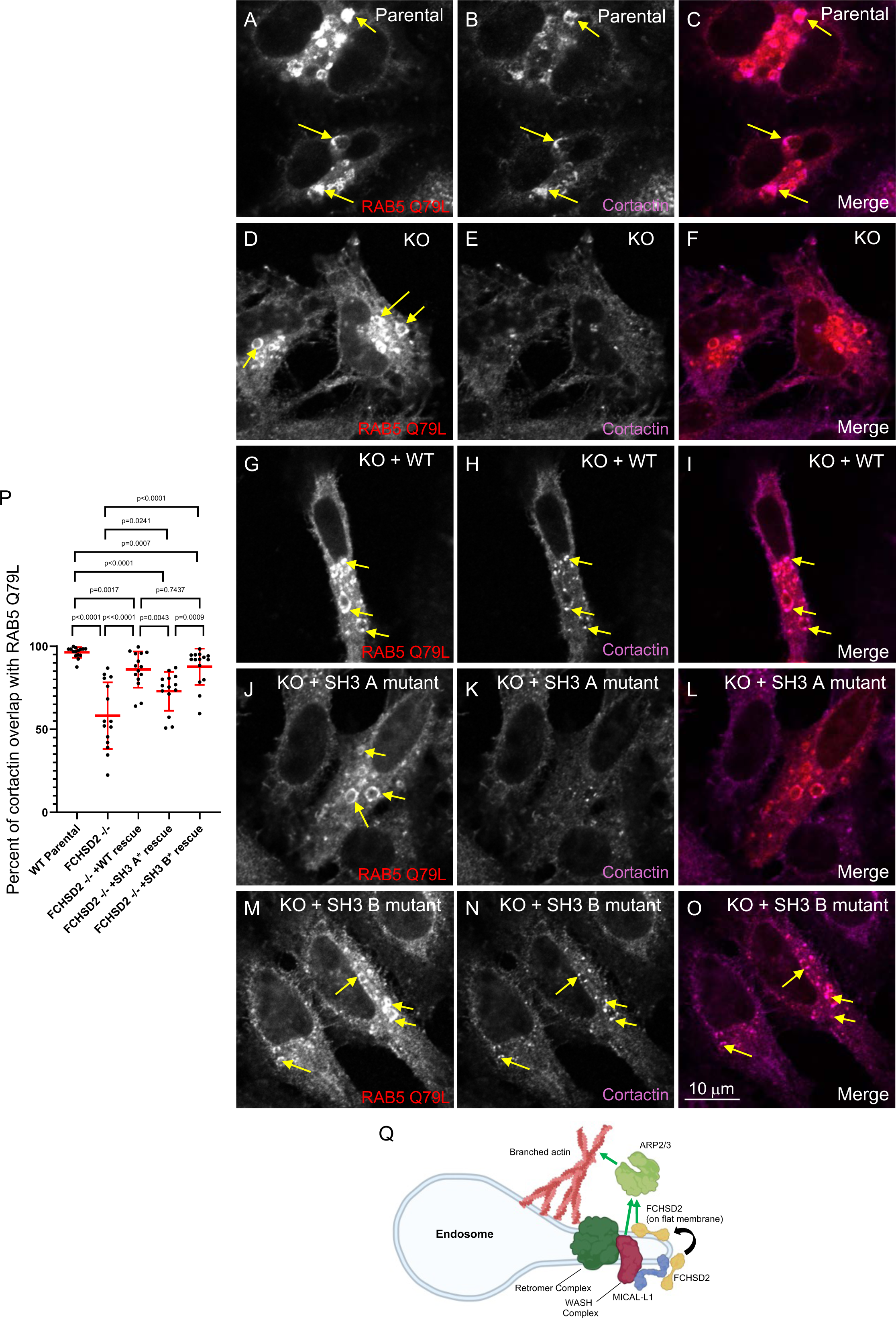
Visualization of decreased branched actin at endosomes in FCHSD2 knock-down cells. A-O. WT parental (A-C), FCHSD2 knock-out (D-F), FCHSD2 knock-out + WT FCHSD2 rescue (G-I), FCHSD2 knock-out + FCHSD2 SH3 A mutant rescue (J-L), and FCHSD2 knock-out + FCHSD2 SH3 B mutant rescue (M-O) were transfected with mCherry-RAB5 Q79L. Cells were fixed and immunostained with an antibody against cortactin to mark branched actin. WT parental cells show a robust cortactin localization at RAB5 Q79L endosomes (B, yellow arrows), whereas the FCHSD2 knock-out cells show a significant decrease in cortactin localized to endosomes (E). Transfection of WT FCHSD2 rescues cortactin localization to the RAB5 endosomes (H, yellow arrows). Transfection with the FCHSD2 SH3 A mutant displays impaired rescue of cortactin at endosomes (K), whereas transfection of FCHSD2 SH3 B mutant rescued cortactin endosomal localization (N, yellow arrows). These data suggest that the FCHSD2 SH3 A domain is required to promote branched actin polymerization at endosomes. P. Quantification of A-O. Confocal Z-stack images were captured and analyzed by Imaris software using the surfaces function. mCherry-RAB5 Q79L structures were 3D rendered and a region of interest around the enlarged endosomes was demarked. Cortactin puncta in this region were also 3D rendered. Cortactin surfaces that contacted RAB5 structures were filtered by setting the maximal “shortest distance to surface” at 1x10^-7^ nm. The volume of cortactin structures that contacted RAB5 structures was summed and represented as a percentage of the total cortactin volume in the designated region of interest. Q. Model for the role of FCHSD2 in fission at endosomes. FCHSD2 is recruited to and/or stabilized at endosomes through an interaction with MICAL-L1. FCHSD2 induces branched actin polymerization to promote endosome fission and receptor recycling.

## Discussion

Current understanding of fission at the endosome has lagged behind knowledge of clathrin-coated pit (CCP) scission at the PM, just as internalization has been more extensively studied than receptor recycling. However, many of the defined steps of CCP scission are analogous to fission at EE/SE, with the actin cytoskeleton playing a central role in the events at both membranes, and both processes will be referred to herein as “fission.” Indeed, actin polymerization increases at clathrin-coated pits at the late stages of CCP internalization (Merrifield *et al*, 2002), in concert with increased dynamin concentrations (Grassart *et al*, 2014). At EE/SE there is a dynamic polarized branched actin network that generates a pushing force on the endosome (Derivery *et al*., 2009). Both CCP fission at the PM and endosome fission are facilitated by the ARP2/3 complex, which nucleates actin and generates branched actin (Pollard, 2007). However, the complex that serves as a nucleation promotion factor at endosomes and activates ARP2/3 is the WASH complex, whereas the N-WASP complex promotes branched actin via ARP2/3 at the PM (Derivery *et al*., 2009; Merrifield *et al*, 2004). An additional distinction between CCP and endosome fission is that the final step of CCP release requires GTP hydrolysis of dynamin, whereas evidence supports involvement of either dynamin (Derivery *et al*., 2009) and/or the dynamin-family ATPase EHD1 in fission at the endosome (Cai *et al*., 2012; Cai *et al*., 2013; Cai *et al*., 2014; Deo *et al*., 2018; Jones *et al*., 2020; Kamerkar *et al*., 2019).

Finally, a key difference between CCP fission and endosomal fission is the involvement of different scaffold proteins in the process. At the PM, intersectin 1 plays an important role in the recruitment of the actin regulator FCHSD2 as well as dynamin (Almeida-Souza *et al*., 2018; Sengar *et al*, 1999), whereas at endosomes, we show that MICAL-L1 recruits FCHSD2 (Figs. 1 and 2) and EHD1 (Sharma *et al*., 2009). Notably, MICAL-L1 has been linked to microtubules (Xie *et al*., 2019) and motor proteins (Rahajeng *et al*, 2010), and the tension between microtubule-based pulling and branched actin pushing on membranes has been proposed as a unique feature in endosomes that causes membrane tension and leads to fission (Roux *et al*, 2006).

MICAL-L1 is a RAB8 effector and key endosomal protein that localizes to both vesicular and tubular endosomes and recruits EHD1 to carry out the final steps of endosome fission (Cai *et al*., 2014; Farmer *et al*., 2020; Giridharan *et al*, 2012; Giridharan *et al*., 2013; Rahajeng *et al*., 2012; Sharma *et al*, 2010; Sharma *et al*., 2009). In addition to its known role in recruiting players involved in the end stages of endosome fission, our new data now supports a role for this scaffold in driving the early, actin constriction of endosomal membranes. Several lines of evidence suggest how MICAL-L1 may regulate the actin cytoskeleton at endosomes. First, MICAL-L1 interacts with Syndapin2/PACSIN2 (Giridharan *et al*., 2013), a member of the Syndapin F-BAR containing proteins that constrict and tubulate membranes and regulate actin organization (Dharmalingam *et al*, 2009; Kessels & Qualmann, 2002; Qualmann & Kelly, 2000; Qualmann *et al*., 1999; Wang *et al*, 2009). Second, recent studies provide evidence that MICAL-L1 interacts with two homologous proteins, CIN85 (also known as SH3-domain kinase binding protein 1) (Havrylov *et al*, 2009; Huttlin *et al*, 2017) and CD2AP (CD2 associated protein) (Huttlin *et al*, 2021), each of which contains a capping protein interaction (CPI) motif that can interact with actin capping protein to promote actin branching (Bruck *et al*, 2006; Cooper & Pollard, 1985; McConnell *et al*, 2020). Most significantly, we demonstrate here that MICAL-L1 interacts with the mammalian Nervous Wreck (Nwk) homolog, FCHSD2, which regulates actin assembly in flies and human cells (Almeida-Souza *et al*., 2018; Del Signore *et al*., 2021; Rodal *et al*., 2008; Stanishneva-Konovalova *et al*., 2016).

FCHSD2 has been implicated in receptor recycling (Xiao *et al*., 2018; Xiao & Schmid, 2020) and Nwk localizes to recycling endosomes (Rodal *et al*., 2011). Indeed, consistent with roles for FCHSD2 in both endocytic function and actin regulation, FCHSD2 regulates ciliogenesis and knock-out mice display acoustic vulnerability and hearing loss (Wang *et al*, 2022; Zhai *et al*, 2022a). Moreover, Nwk activates WASP to promote ARP2/3-dependent actin filament assembly (Stanishneva-Konovalova *et al*., 2016), and similarly FCHSD2 stimulates ARP2/3 on flat membranes (Almeida-Souza *et al*., 2018). FCHSD2 and CDC42 can also simultaneously bind N-WASP, providing an additional layer of actin regulation (Rodal *et al*., 2008; Zhai *et al*., 2022b). Given that the WASH complex is the major activator of endosomal ARP2/3 actin nucleation and WASH function is required for fission and receptor recycling (Derivery *et al*, 2012; Derivery *et al*., 2009), it is logical to speculate that FCHSD2 activates WASH at endosomes to promote ARP2/3-based actin nucleation. Our experiments do not distinguish whether the requirement for the FCHSD2 SH3 A domain in endosomal actin assembly is due to MICAL-L1 interactions, actin assembly through WASP family proteins, or both. However, we note that FCHSD2 is a dimer, and SH3A valency *in vivo* is likely even higher due to interactions with other binding partners. Therefore, FCHSD2 complexes on endosomes are likely capable of interacting with both MICAL-L1 and WASP family proteins. In addition, affinity capture-mass spectrometry has identified FCHSD2 as a SNX27 interactor (Shi *et al*, 2021), a protein that binds directly to both the WASH complex and the retromer (Steinberg *et al*, 2013; Temkin *et al*., 2011), further supporting endosomal WASH complex activation. Indeed, our experiments show that FCHSD2 knock-out cells have impaired branched actin generation at endosomes (Fig. 8 and EV5), and that inhibition of ARP2/3 function leads to a failure to generate branched actin networks on endosomes (EV4). Overall, these findings support a potential role for MICAL-L1 as a bridge or link between early actin-based steps of endosomal membrane constriction via FCHSD2 and EHD1-mediated fission of endosomes.

One intriguing question is what regulates the recruitment of FCHSD2 to endosomes? Nwk and FCHSD2 appear to be autoinhibited by both SH3 domains, although the SH3 B domain plays a more predominant role (Almeida-Souza *et al*., 2018; Kelley *et al*, 2015). It has been proposed that intersectin 1 recruits FCHSD2 to clathrin coats as they mature, by an atypical SH3-SH3 interaction with the SH3 B domain of FCHSD2, while keeping FCHSD2 in a state of low activation (Almeida-Souza *et al*., 2018). Subsequently, PI(3,4)P2 accumulation at the edge of the CCP allows BAR domain binding and recruitment to the flat membrane surrounding the CCP. This in turn leads to the activation of FCHSD2 and induction of actin polymerization. At endosomes, however, MICAL-L1 recruitment of FCHSD2 is likely via one of the ∼14 MICAL-L1 proline rich domains with the SH3 A domain of FCHSD2 (Fig. 1). We envision that analogous to the process at the PM, the BAR domain of FCHSD2 then interacts with endosomal phospholipids and is stabilized on the flat membrane region of endosomes to promote of branched actin generation. Indeed, FCHSD2 can bind to phosphatidylinositol(3,4,5)-trisphosphate (Almeida-Souza *et al*., 2018) which is generated on endosomes by class I phosphatidylinositol-3-kinase activity (Jethwa *et al*, 2015).

It is unclear at present whether FCHSD2 is “handed-off” from CCP at the PM to endosomes, or whether it is recruited from a cytoplasmic pool. However, recent studies have demonstrated that stimulating receptor-mediated endocytosis leads to a significant increase in both the number and size of EE/SE (Naslavsky & Caplan, 2023b), as well as recruitment of EHD1 and fission machinery from the cytoplasm to endosomes (Dhawan *et al*., 2020). As a result, this may lead to an increased level of MICAL-L1 available for the recruitment of FCHSD2. It is possible that FCHSD2 is recruited from CCP at the PM. In this scenario, the SH3-SH3 interactions between FCHSD2 and intersectin 1 may be higher affinity than the FCHSD2 SH3 A domain with MICAL-L1 proline rich domains, but by increasing the availability of the MICAL-L1 binding sites for FCHSD2, increased recruitment of the latter protein may occur. Another possibility is that intersectin 1 itself is initially involved in FCHSD2 recruitment to endosomes. Indeed, a shortened form of intersectin (ITSN-s) serves as an effector for RAB13 (Ioannou *et al*, 2017), ITSN-1 interacts with endosomal components such as RAB5 and ARF6 (Wong *et al*, 2012), and ITSN-2 has been implicated in the regulation of endosomal recycling (Gubar *et al*, 2020). However, the relationship between intersectins, MICAL-L1 and FCHSD2 at endosomes remains to be elucidated.

Our data is consistent with a model in which MICAL-L1 is initially required for recruitment of FCHSD2 to endosomal membranes (see model; Fig. 8Q). This interaction occurs via the SH3 A domain of FCHSD2, and likely requires one of the multiple MICAL-L1 proline-rich domains domains. The identification of the specific proline rich region(s) remains unknown, and it is possible that several such regions may be capable of binding. Once binding to MICAL-L1 occurs, we speculate that FCHSD2 migrates to extended regions of the budding endosomal membrane, consistent with its preference for flat membranes and select phosphoinositides (Almeida-Souza *et al*., 2018). As a result, this membrane association helps stimulate WASH-mediated ARP2/3 activation of branched actin, leading to further membrane constriction and tubulation, supported by RAB and motor proteins (Farmer *et al*., 2020). Ultimately, Coronin 1C (Hoyer *et al*., 2018) and 2A (Dhawan *et al*., 2022) have been implicated in clearance of branched actin, providing accessibility for the MICAL-L1 partner and fission protein, EHD1, to the membrane, leading to vesicle/tubule release and cargo recycling to the PM. Overall, our study provides new insight into the mechanisms of EE/SE fission, and helps identify how actin-based constriction events are coupled with nucleotide hydrolysis to promote fission.

## Materials and Methods

### Antibodies and reagents

The following antibodies were used: anti-MICAL-L1 (1794, LifeTein, 1:200 for immunoblotting), anti-GST-HRP (A01380, Genscript, 1:500), anti-GAPDH-HRP (HRP-60004, Proteintech, 1:5000), anti-FLAG (F1804, Sigma, 1:800), anti-FCHSD2 (described in (Almeida-Souza *et al*., 2018), 1:300 for immunostaining), anti-FCHSD2 (PA5-58432, Invitrogen, 1:250 for immunoblotting), anti-MHC-1 (purified W6/32, Leinco Technologies), anti-EEA1 (3288, Cell Signaling, 1:30), anti-cortactin (05-180-I, Sigma, 1:200), anti-acetylated tubulin (3971, Cell Signaling Technology, 1:300), anti-CP110 (12780, Proteintech, 1:200), anti-MICAL-L1 (H00085377-B01P, Novus, 1:500 for immunostaining), anti-MICAL-L1 (ab220648, Abcam, 1:300 for immunostaining), donkey anti-mouse-HRP (715-035-151, Jackson, 1:5000), mouse anti-rabbit IgG light chain-HRP (211-032-171, Jackson, 1:3000), Alexa Fluor 568–conjugated goat anti-rabbit (A11036, Molecular Probes, 1:500), Alexa Fluor 568–conjugated goat anti-mouse (A21043, Molecular Probes, 1:500), Alexa Fluor 488–conjugated goat anti-rabbit (A11034, Molecular Probes, 1:500), Alexa Fluor 488–conjugated goat anti-mouse (A11029, Molecular Probes, 1:500), and Alexa Fluor 647-conjugated goat anti-mouse (115-606-008, Jackson ImmunoResearch, 1:750). The following plasmid constructs were used: GFP-RAB5 Q79L (Roberts *et al*, 1999), mCherry-RAB5 Q79L (35138, Addgene), FLAG-FCHSD2 (GenScript), FCHSD2-GFP (Almeida-Souza *et al*., 2018), FCHSD2 Y576S+F607S – GFP (Almeida-Souza *et al*., 2018), and FCHSD2 YY478/480AA-GFP (Almeida-Souza *et al*., 2018). The following reagents were used: Sepharose resin (L00206, GenScript), Alexa fluor 488– conjugated transferrin (T13342, Invitrogen), CF-568–conjugated Phalloidin (44-T VWR, Biotium, 1:100), PI3K inhibitor LY294002 (501099125, Fisher Scientific), CK-689 (182517, MilliporeSigma), and CK-666 (182515, MilliporeSigma). Yeast two-hybrid screens of over 100,000,000 potential interactions were performed by Hybrigenics (Boston, MA) using the full length wild-type MICAL-L1 as bait.

### Cell culture and treatments

The HeLa cell line (ATCC-CCL-2) was obtained from ATCC and cultured with complete DMEM (high glucose) (ThermoFisher Scientific, Carlsbad, CA) with 10% fetal bovine serum (FBS) (Sigma-Aldrich), 1× penicillin-streptomycin, and 2 mM L-glutamine. The hTERT RPE-1 human epithelial cell line (ATCC-CRL4000) was obtained from ATCC and grown in DMEM/F12 (ThermoFisher Scientific, Carlsbad, CA) with 10% fetal bovine serum (FBS), 1× penicillin-streptomycin, 2 mM L-glutamine, and 1X non-essential amino acids (ThermoFisher Scientific, Waltham, MA). The non-small cell lung cancer cell (NSCLC) line H-1650 was obtained from ATCC (CRL-5883) and cultured in RPMI with 10% FBS, 1× penicillin-streptomycin, 2 mM L-glutamine, 1X MEM non-essential amino acids, 25 mM HEPES, and 1mM sodium pyruvate. Validated CRISPR/Cas9 gene-edited HeLa knock-out cells (FCHSD2^-/-^ and MICAL-L1^-/-^) were obtained from GenScript (Piscataway, NJ). All media also contained 100 μg/ml Normocin (Invitrogen) to prevent mycoplasma and other contamination and cells were routinely tested for mycoplasma contamination. All cells were cultured at 37°C in 5% CO_2_. The small interfering siRNA (siRNA) oligonucleotide targeting human FCHSD2 (5’-GCAUACUCCUGAGACCUCA[dT][dT]-3’) was obtained from Sigma Aldrich. FCHSD2 siRNA knockdown in all HeLa cell lines was performed for 48 h using the DharmaFECT transfection reagent (Dharmacon, Lafayette, CO). To achieve knockdown in both RPE and NSCLC cells, siRNA was transfected using the Lipofectamine RNAi/MAX (Invitrogen, Carlsbad, CA) reagent for 72 h. Knockdown efficiency was confirmed via immunoblotting. Transfection with FCHSD2 constructs and RAB5 Q79L constructs was achieved using the FuGENE 6 (Promega, Madison, WI) transfection reagents and protocol. The DNA ratios used for co-transfections are noted below.

### Immunoblotting

Cultured cells were washed three times with ice-cold PBS and harvested with a cell scraper. Pelleted cells were resuspended in lysis buffer (50 mM Tris–HCl pH 7.4, 150 mM NaCl, 1% NP-40, 0.5% sodium deoxycholate) with freshly added protease inhibitor cocktail (Roche, Indianapolis, IN) for 30 min on ice. Lysates were then centrifuged at 13,000 rpm at 4°C for 10 min. Following centrifugation, 4x loading buffer was added to each sample and then boiled for 10 min. Collected lysates or samples from the GST pulldowns were separated by 10% SDS-PAGE, and transferred onto a nitrocellulose membrane (GE Healthcare, Chicago, IL). After membrane transfer, the membranes were blocked with 5% dried milk in PBS containing 0.3% (v/v) Tween-20 (PBST) for 30 min at room temperature. The membrane was incubated with the primary antibody diluted in PBST at 4°C overnight. The membrane was washed three times with PBST after incubation with the primary antibody and then incubated at room temperature for 30 min with the appropriate HRP-conjugated secondary antibody diluted in PBST. Enhanced chemiluminescence (Bio-RAD, Hercules, CA) was used to visualize the HRP-conjugated secondary antibody and digital images were acquired with iBright Imaging Systems (Invitrogen).

### Recombinant gene expression and protein purification

The GST-tagged FCHSD2 DNA constructs were transformed into *Escherichia coli* Rosetta strain, and freshly transformed *E. coli* was inoculated in 50 ml of Luria-Bertani (LB) broth, containing 100 μg/ml ampicillin. Following an overnight incubation at 37°C with shaking, the 50 ml culture was used to inoculate a 1000 ml LB culture (containing antibiotic), which was incubated at 37°C with shaking until the OD reached 0.4-0.6 at 600 nm. Protein expression was then induced with 1 mM IPTG, and the culture was incubated overnight at 18°C. The bacteria were then centrifuged at 5000 rpm for 15 min at 4°C. The bacterial pellet was then resuspended in ice-cold lysis buffer composed of 1x PBS (pH 7.4) and 1 tablet of protease inhibitor (Roche) per 10 ml. Following resuspension, the sample was lysed on ice via sonication (10 min of total sonication with 20 s on and 10 s off cycles). To separate the cellular debris, lysates were then centrifuged at 19,000 rpm for 45 min at 4°C. The supernatant was then incubated with Glutathione Sepharose resin overnight at 4°C. Three subsequent washes were performed with 10-bed volumes of wash buffer, which was composed of 1x PBS with protease inhibitor.

### GST-pulldown

Following protein purification, 20 µg of bead-bound GST constructs were centrifuged at 13,000 rpm for 30 s and resuspended in 30 µl of 1x PBS. 1 U micrococcal nuclease was added to the bead-bound protein and incubated at 30°C for 10 min. Following this incubation, HeLa cell lysate (lysed with 1% Brij98, 25 mM Tris-HCl, 125 mM NaCl, 1 mM MgCl_2_, protease inhibitor, pH 7.4) was added to the bead-bound protein and incubated at 4°C for 3 h. The samples were then washed 3 times with 10-bed volumes of wash buffer (0.1% Brij98, 25 mM Tris-HCl, 125 mM NaCl, 1 mM MgCl_2_, protease inhibitor, pH 7.4) and eluted with 4x loading buffer before undergoing SDS-PAGE and immunoblotting.

### FCHSD2 localization at RAB5 endosomes

HeLa cells were plated on coverslips and co-transfected with GFP-FCHSD2 and mCherry-RAB5 Q79L (1:1) using the FuGene transfection system. Following overnight transfection, coverslips were fixed with 4% paraformaldehyde (in PBS) for 20 min at room temperature. After fixation, the coverslips were washed 3x with PBS and mounted in Fluoromount (ThermoFisher Scientific). Confocal images were captured using a Zeiss LSM 800 confocal microscope (Carl Zeiss) with a 63×/1.4 NA oil objective, and the images were analyzed in ImageJ (National Institutes of Health, Bethesda, MD). In ImageJ, a profile 2.81 µM in length was drawn from the cytoplasm into the enlarged mCherry-RAB5 Q79L vesicles. The fluorescence intensities along the profile were collected for both the red and the green channels. Background subtraction was performed using the background fluorescence intensity in the region of the cytoplasm devoid of mCherry-RAB5 Q79L vesicles. Following background subtraction, fluorescent intensities along the profile were calculated as relative intensities of the maximal intensity for both the red and the green channels. The distance from the maximal mCherry-RAB5 Q79L fluorescent intensity was normalized for each figure and plotted. Localization of endogenous FCHSD2 at GFP-RAB5 Q79L structures in WT parental and MICAL-L1 knockout cells was quantified using Imaris 9.9.1 software. A region of interest (ROI) containing the RAB5 vesicles was marked, and all FCHSD2 puncta and RAB5 Q79L structures in that area were three-dimensionally (3D)-rendered (see parameters: Table 1, Rows 1 and 2). The volumetric sum of all FCHSD2 puncta that contacted a RAB5 Q79L structure was calculated as a percentage of the entire FCHSD2 volume in that ROI.

**Table 1.** Imaris parameters table.

| Reference number | Description | Surface grain size (μm) | Diameter of largest sphere (μm) | Manual threshold value | Region growing estimated diameter (μm) | Quality above |
| --- | --- | --- | --- | --- | --- | --- |
| 1 | **FCHSD2 localization:**<br>FCHSD2 puncta | 0.0800 | 0.200 | 740.43 | 0.600 | -0.181 |
| 2 | **FCHSD2 localization:**<br>Rab5 QL structures | 0.0500 | 0.400 | 5194.76 | 0.500 | 3962 |
| 3 | **Endosome morphology:** EEA1 | 0.0500 | 0.400 | 4073.89 | 0.500 | 6679 |
| 4 | **Endosome morphology:** MICAL-L1 | 0.198 | 1.00 | 5751.4 | N/A | N/A |
| 5 | **Fission assay:** Tf and EEA1 | 0.250 | 3.48 | 5746 | 1.30 | 427 |
| 6 | **Cortactin at Rab5 QL:** mCherry-Rab5 QL | 0.0500 | 1.00 | 9696.54 | 0.500 | 5148 |
| 7 | **Cortactin at Rab5 QL:** GFP-Rab5 QL | 0.0500 | 0.400 | 5194.76 | 0.500 | 3962 |
| 8 | **Cortactin at Rab5 QL:** Cortactin | 0.0800 | 0.200 | 3388.84 | 0.600 | -0.128 |
| 9 | **EEA1-Cortactin contacts:** EEA1 | 0.200 | 0.638 | 2.69884 | N/A | N/A |
| 10 | **EEA1-Cortactin contacts:** cortactin | 0.170 | 0.543 | 2.46539 | N/A | N/A |

### Recycling assays

The transferrin recycling assay was performed by diluting Transferrin-Alexa Fluor 488 (Tf-488; Invitrogen) 1:700 in complete DMEM. FCHSD2 siRNA- or Mock-treated cells were allowed to uptake transferrin for 10 min at 37°C. Following uptake, cells were washed twice with 1x PBS and then chased with complete DMEM for 50 min. After three washes with 1x PBS, cells were fixed with 4% paraformaldehyde in PBS for 20 min and then mounted in Fluoromount. For the MHC-1 recycling assay, cells were incubated with anti-MHC-1 antibody diluted in complete DMEM for 20 min at 37°C. After uptake, the remaining antibodies bound to the cell surface MHC-1 were removed via a 1 min glycine strip (0.1 M HCl-glycine, pH 2.7). The glycine-stripped cells were then either washed and fixed (uptake group) or washed 3 times with 1x PBS and chased with complete DMEM for 50 min at 37°C. The cells were then washed 3 times with PBS and fixed with 4% paraformaldehyde in PBS for 20 min. Cells were then stained with fluorochrome–conjugated secondary antibodies for 30 min, washed 3 times with PBS, and mounted in Fluoromount. Confocal images of cells after both the transferrin and MHC-1 recycling assays were captured using a Zeiss LSM 800 confocal microscope (Carl Zeiss) with a 63×/1.4 NA oil objective. The arithmetic mean of fluorescence intensity for each image was analyzed using Zen Blue software. The fluorescence intensity after the chase was represented as a percentage of the fluorescent intensity after uptake and plotted as “percent remaining in cells.”

### Measurement of endosome size

FCHSD2 siRNA- or Mock-treated HeLa cells were fixed with 4% paraformaldehyde (PBS) for 20 min at room temperature. After fixation, the cells were stained with anti-EEA1 for 1 h and the appropriate fluorochrome–conjugated secondary antibody for 30 min at room temperature. The coverslips were mounted, and images were captured using a Zeiss LSM 800 confocal microscope (Carl Zeiss) with a 63×/1.4 NA oil objective. EEA1 area was measured using Imaris 9.9.1 by rendering surfaces according to the settings listed on Table 1, Row 3. For measuring MICAL-L1 endosome area, FCHSD2 knock-out cells were transfected with Flag-FCHSD2 as the rescue group. WT parental, FCHSD2 knock-out, knock-out + rescue, mock-treated, and FCHSD2 siRNA-treated cells were fixed (as described above) and stained with anti-MICAL-L1. The MICAL-L1 area per cell was measured using Imaris 9.9.1 with settings according to Table 1, Row 4.

### Fission assay

Mock- and FCHSD2 siRNA-treated HeLa cells were incubated for 45 min in DMEM containing 80 μM of the PI3K inhibitor LY294002 (Cayman Chemical, Ann Arbor, MI) diluted for 1 h at 37°C. Following the initial incubation, cells were washed with PBS and then incubated in DMEM containing Transferrin-Alexa Fluor 488 (Invitrogen) and 80 μM LY294002 for 15 min at 37°C. Cells were then washed with PBS to remove the inhibitor and chased with complete DMEM for 20 min to allow synchronized fission and recycling. Finally, cells were washed 3 times with PBS and fixed with 4% paraformaldehyde in PBS for 20 min at room temperature. Cells on coverslips were incubated with anti-EEA1 for 1 h, followed by 3 PBS washes and a 30 min incubation with the appropriate 568 fluorochrome–conjugated secondary antibody. After staining, coverslips were mounted in Fluoromount, and images were captured using a Zeiss LSM 800 confocal microscope (Carl Zeiss) with a 63×/1.4 NA oil objective. Quantification of the assay was done using surface rendering in Imaris 9.9.1. Tf-containing vesicles and EEA1- decorated endosomes were 3D rendered according to the parameters on Table 1, Row 5. The size of Tf-containing vesicles was plotted in GraphPad Prism as a frequency distribution (interleaved) graph with bins from 2 μm to 10 μm and the bin width set at 1 μm. A Gaussian curve was plotted to infinity on the frequency distribution graphs for all four experimental groups (mock uptake, mock chase, knock-down uptake, knock-down chase). The area under each of the curves was calculated using the “Area under the curve” function in GraphPad Prism 10.2.3. The area between the curves for the mock- or siRNA-treated group was calculated as the area under the curve after uptake minus the area under the curve after chase.

### Quantification of cortactin at RAB5 QL endosomes

Experimental groups included HeLa WT parental, FCHSD2 knock-out cells, and FCHSD2 knock-out cells transfected with FCHSD2-GFP, FCHSD2 YY478/480AA-GFP, or FCHSD2 Y576S+F607S-GFP. Cells were transfected with mCherry-RAB5 Q79L (Addgene #35138; 1:1 in cells with co-transfection) overnight. GFP-RAB5 Q79L was transfected into WT parental, FCHSD2 knock-out, and MICAL-L1 knock-out cells. Cells were washed with PBS and fixed with 4% paraformaldehyde in PBS for 20 min at room temperature. Following fixation, cells were stained with anti-cortactin for 1 h, followed by 3 washes with PBS and a 30 min incubation with the appropriate 647 fluorochrome–conjugated secondary antibody. Three additional washes with PBS were performed after secondary incubation, and then the cells were mounted in Fluoromount. Z-stacks were captured using a Zeiss LSM 800 confocal microscope (Carl Zeiss) with a 63×/1.4 NA oil objective. Using Imaris 9.9.1 software, mCherry-RAB5 Q79L and GFP-RAB5 Q79L structures were 3D-rendered (see Table 1, Rows 6 and 7, respectively). A region of interest around the active Rab5 structures was demarked, and cortactin structures in this region were 3D-rendered (see Table 1, Row 8). The cortactin structures that contacted the active RAB5 structures were selected using the filter tab and setting the maximal “shortest distance to surface” at 1x10^-7^ nm. The volume of cortactin structures that contacted RAB5 structures was summed and represented as a percentage of the total cortactin volume in the designated region of interest.

### Actin inhibitors

HeLa cells were plated on coverslips and transfected with GFP-RAB5 Q79L. The following day, the cells were incubated for 40 min with 300 μM of either the control CK-689 or the specific ARP2/3 inhibitor CK-666 diluted in complete DMEM. Following the incubation, the cells were washed with PBS and fixed with 4% paraformaldehyde in PBS for 20 min. The cells were then washed 3 times with PBS and stained with 568-phalloidin for 1 h to mark the filamentous actin network. The cells were washed an additional 3 times and then mounted on slides in Fluoromount. Confocal images were captured using a Zeiss LSM 800 confocal microscope (Carl Zeiss) with a 63×/1.4 NA oil objective.

### EEA1-Cortactin contacts

Mock- and FCHSD2 siRNA-treated NSCLC cells were washed with PBS and fixed with 4% paraformaldehyde in PBS for 20 min. The cells were co-stained with anti-EEA1 and anti-cortactin antibodies by incubating with the primary antibody at room temperature for 1 h. The cells were subsequently washed 3 times with PBS and incubated with the appropriate fluorochrome–conjugated secondary antibodies. Following another 3 washes with PBS, cells were mounted in Fluoromount, and images were captured using a Zeiss LSM 800 confocal microscope (Carl Zeiss) with a 63×/1.4 NA oil objective. Images were quantified in the Imaris 9.9.1 software using the surfaces function with the respective parameters for EEA1 (Table 1, Row 9) and cortactin (Table 1, Row 10). Three regions of interest were demarked in the periphery of the cells (defined as an area with no border closer than 5 μm to the nucleus), and another 3 regions in the perinuclear area (defined as an area having a border within 1 μm of the nucleus). EEA1 structures contacting cortactin were defined as EEA1 surfaces with a shortest distance to a cortactin surface with a value of zero. The EEA1 structures that contacted cortactin were represented as a percentage of the total number of EEA1 structures.

### Ciliogenesis assay

To induce ciliogenesis, Mock- and FCHSD2 siRNA-treated RPE-1 cells were shifted to starvation media (DMEM/F12, 0.2% FBS, 1% penicillin-streptomycin, 2 mM L-Glutamine, 1X non-essential amino acids) for 4 h at 37°C in 5% CO_2_. Cells were then rinsed once with cold PBS and fixed with 100% methanol at -20°C for 5 min. After fixation, coverslips were washed 3 times with PBS, and a pre-staining incubation was performed for 30 min (0.5% Triton X and 0.5% BSA in PBS). Coverslips were co-stained with primary anti-CP110 and anti-acetylated-tubulin antibodies for 1 h at room temperature. The coverslips were subsequently washed 3 times and stained with the appropriate fluorochrome–conjugated secondary antibodies. Following secondary incubation, the coverslips were washed 3 times with PBS and mounted in Fluoromount. Images were taken using a Zeiss LSM 800 confocal microscope (Carl Zeiss) with a 63×/1.4 NA oil objective, and maximal intensity orthogonal projections were obtained and quantified in the Zen Blue software. The number of ciliated and non-ciliated cells was manually counted. Ciliated cells were considered to have an elongated acetylated tubulin stain along with retention of CP110 on one of the two centrioles. Non-ciliated cells typically retained CP110 on both centrioles. The percentage of ciliated cells from the total number of cells was quantified for each image and plotted.

### Statistical analysis

Data for all experiments was collected from 3 independent experiments and graphed with the mean and standard deviation. Normality was determined with a D’Agostino and Pearson (or Shapiro-Wilk) normality test. If the normal distribution assumption was met, an unpaired two-tailed *t-*test was used to assess p-values and determine statistical significance between two groups (p<0.05). In the event that the distribution did not meet the assumption of normality, a Mann-Whitney non-parametric two-tailed test was used to assess significance. All the graphical and statistical tests were done using GraphPad Prism 10.2.3.

## Data Availability

No data from this manuscript requires deposition in a public database.

## Acknowledgments

This work was supported by NIH grant R35GM144102 from the National Institute of General Medical Sciences (SC), and by NIH T32 training grant CA009476 (Cancer Biology Training Program; DF), and by the Research Council of Finland (Research Fellow; LA-S).

## Author Contributions

DF carried out the experimentation and collected and interpreted data, and wrote parts of the manuscript. KD, ABM and BA each contributed to experimentation and generation of specific figures. NN supervised and was involved in data interpretation and experimental design, helped with figure preparation and edited parts of the manuscript. LA-S generated reagents, provided scientific input and shared in editing of the manuscript. SC was responsible for the research overall, obtained funding, laid out the overall research goals, supervised and was involved in data interpretation, figure preparation and writing and editing of the manuscript.

## Disclosure and competing interests statement

The authors have no competing interests.

---

**Expanded View 1.**
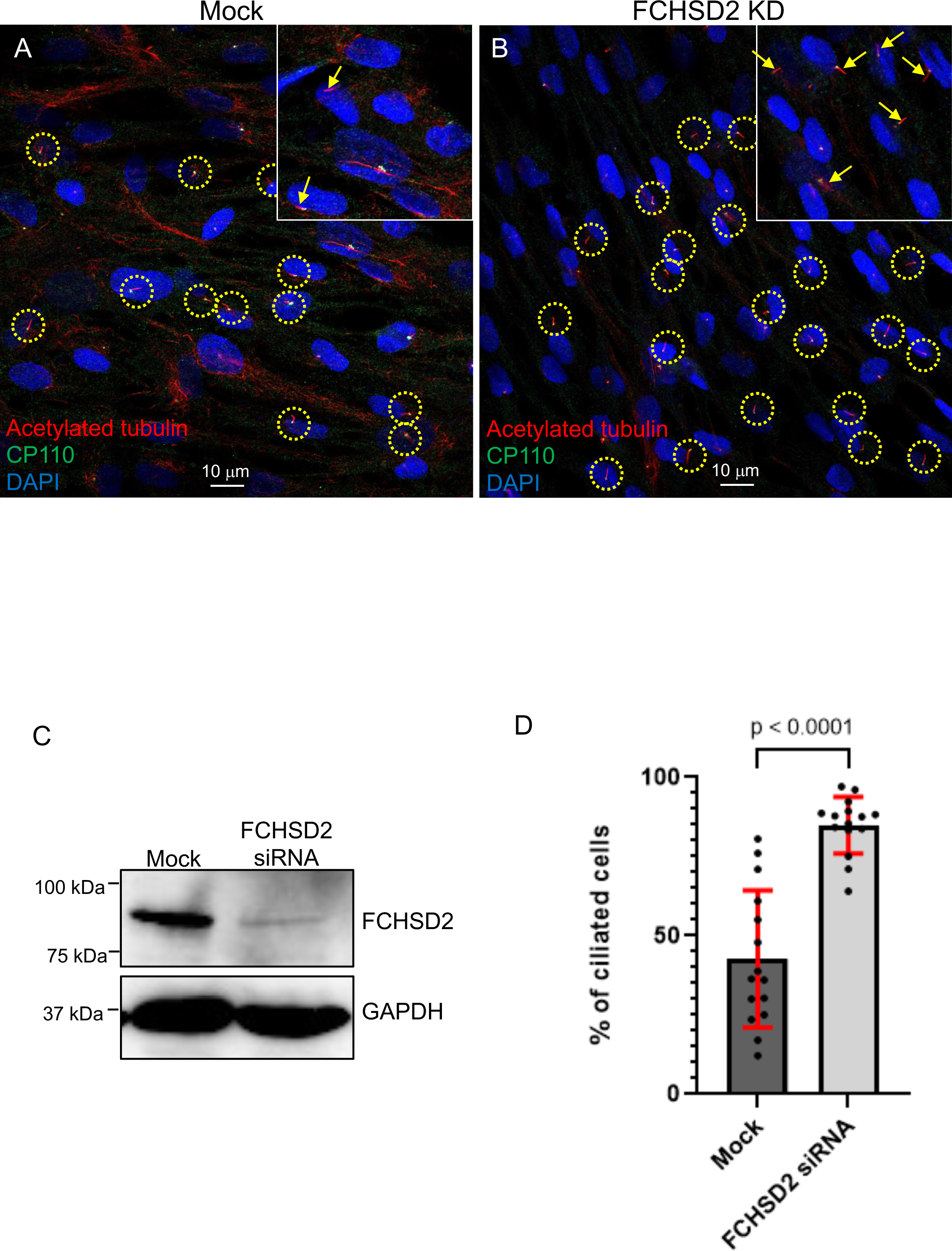
FCHSD2 knockdown increases ciliogenesis. A,B. Mock- or FCHSD2 siRNA-treated RPE-1 cells were serum-starved for 4 h to induce ciliogenesis. Cells were fixed and immunostained with primary antibodies against CP110 (green), acetylated-tubulin (red), and DAPI (blue). C. Immunoblot validation of FCHSD2 siRNA knockdown in RPE cells. D. Percentage of ciliated cells quantified from A,B. The percentage of ciliated cells was manually evaluated by counting the number of ciliated cells and non-ciliated cells in a given field. Ciliated cells were considered to have an elongated acetylated tubulin stain along with retention of CP110 on one of the two centrioles. Non-ciliated cells retained CP110 on both centrioles. The percentage of ciliated cells increases upon FCHSD2 knock-down.

**Expanded View 2.**
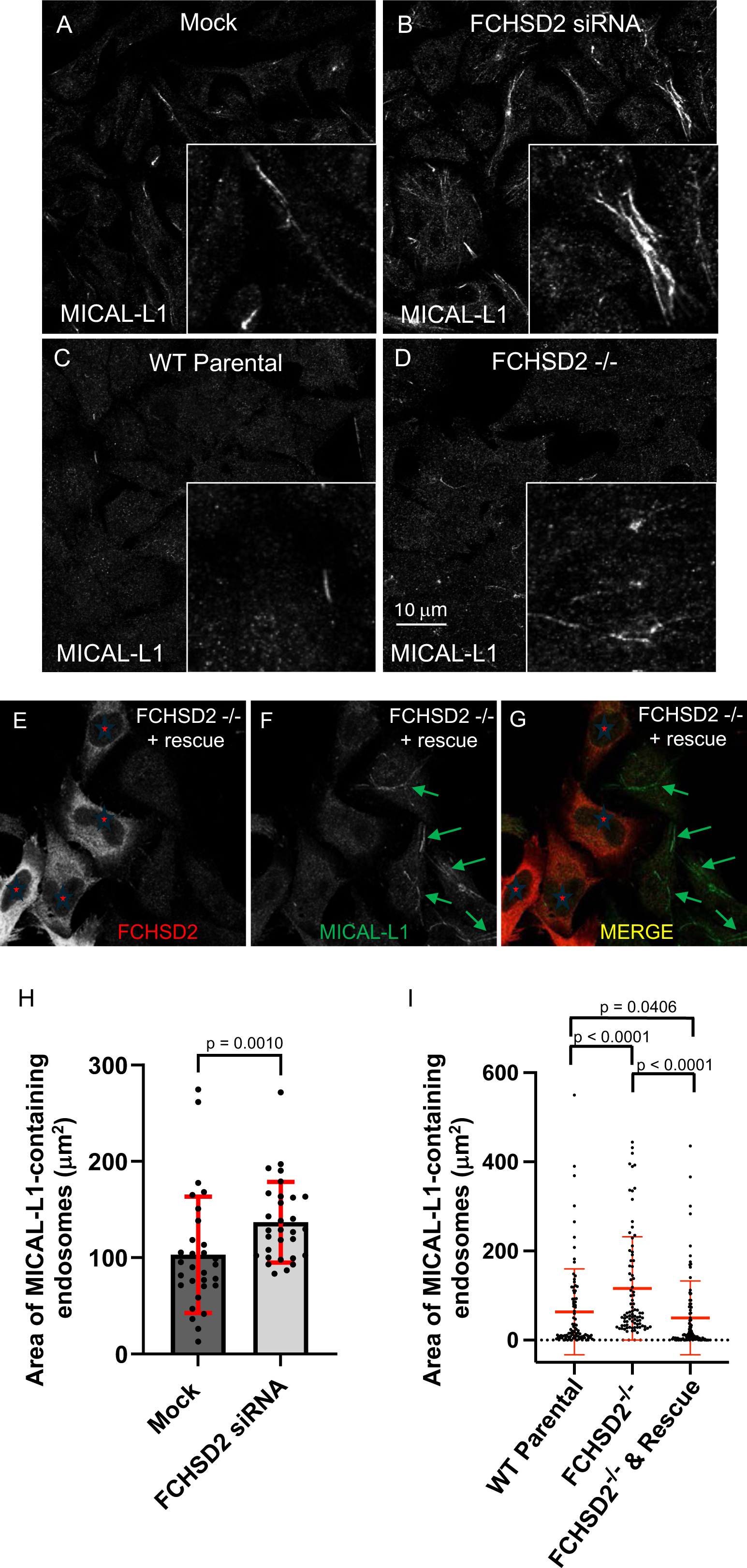
The area of MICAL-L1-decorated tubular recycling endosomes increases upon FCHSD2 knock-down. A-D. Mock, FCHSD2 siRNA, WT parental, and FCHSD2 knock-out HeLa cells were fixed and immunostained with anti-MICAL-L1. Upon knock-down or knock-out, the number and area of MICAL-L1-decorated endosomes increased. E-G. FCHSD2 knockout cells were transfected with FLAG-FCHSD2 and immunostained with anti-FLAG (red) and anti-MICAL-L1 (green). Rescued cells (red star) lack elongated MICAL-L1 endosomes, whereas non-rescued, knock-out cells display an increased number and area of MICAL-L1-decorated endosomes. H. Quantification of increased MICAL-L1 area upon FCHSD2 depletion (A-D). Images were captured and analyzed using the Imaris software by rendering MICAL-L1 surfaces and plotting the MICAL-L1 area per cell. I. Quantification of MICAL-L1 area in WT parental, knock-out, and FCHSD2 rescue cells (E-G). Images were captured and analyzed using the Imaris software. The area of MICAL-L1 surfaces per cell was measured in Imaris and plotted. Validation of knock-down and knock-out is shown in Fig. 5.

**Expanded view 3.**
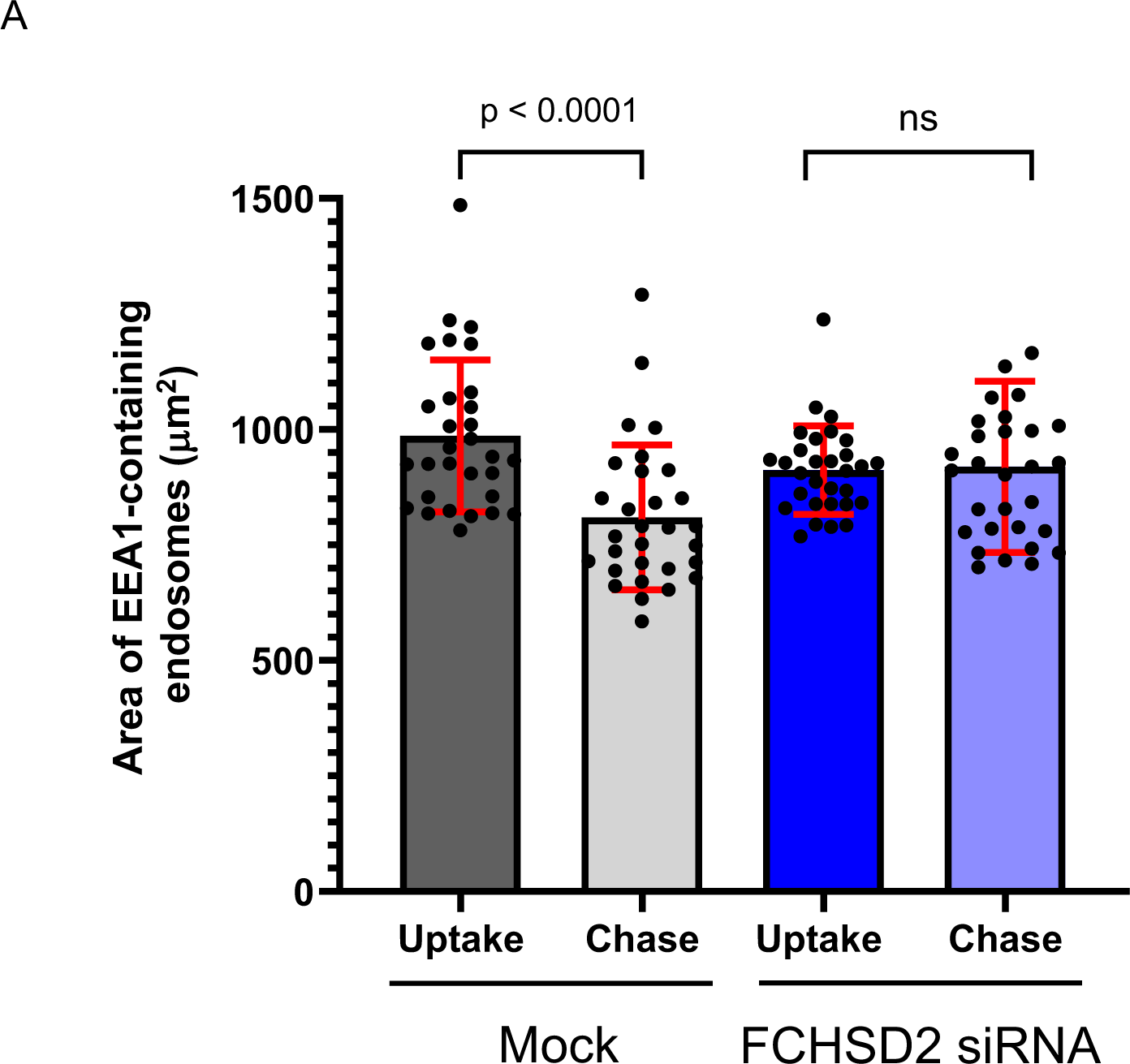
FCHSD2 knockdown leads to impaired fission of EEA1-decorated endosomes. A. Cover-slips from Fig. 6 were co-stained with EEA1 to assess differences in endosome area before and after washout of the PI3K inhibitor. EEA1 surfaces were rendered in Imaris from confocal images. The total EEA1 area per cell was quantified after uptake with the inhibitor and after washout of the inhibitor (chase) for both the mock- and FCHSD2 siRNA-treated cells.

**Expanded view 4.**
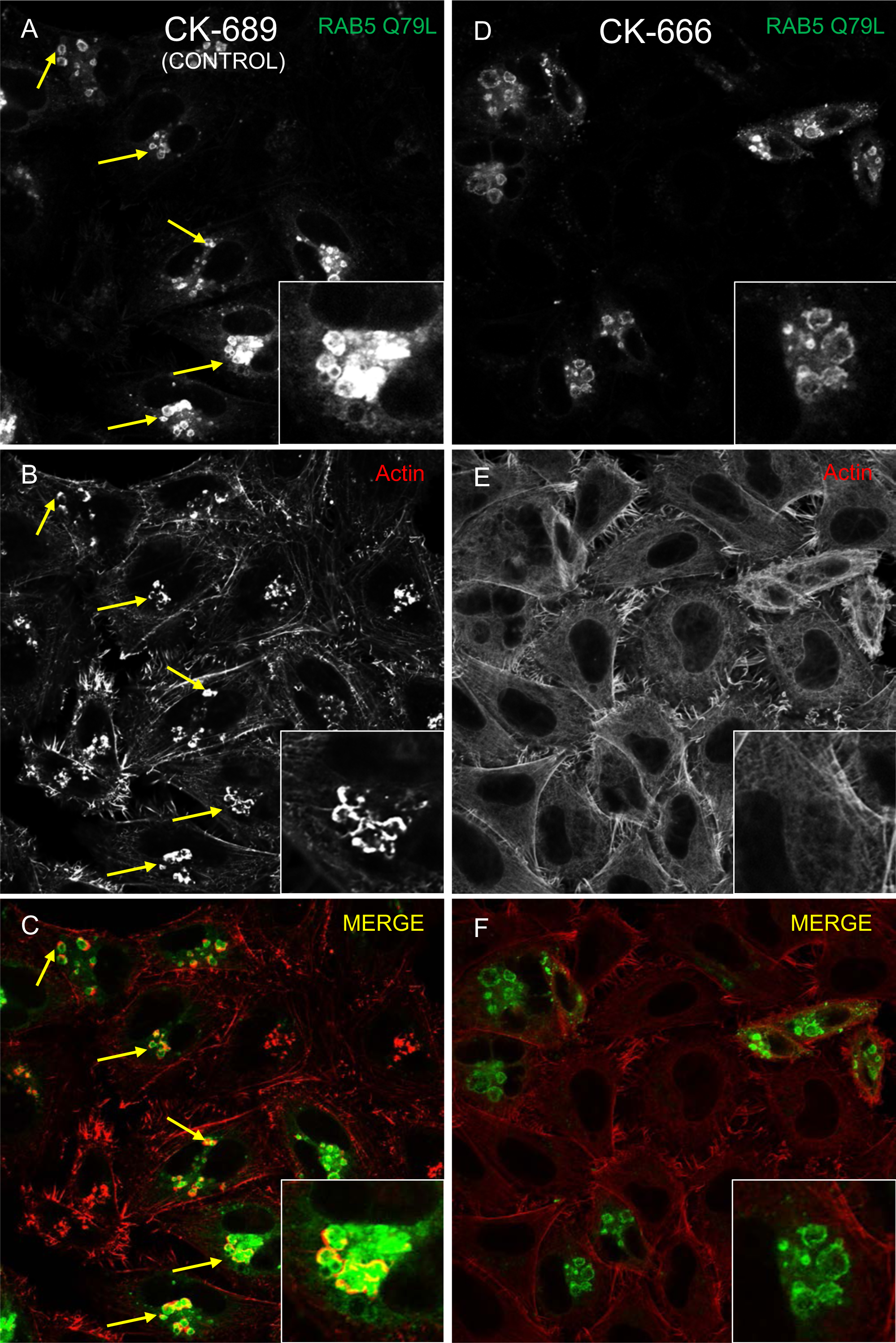
Actin polymerization at RAB5 Q79L endosomes requires ARP2/3. A-F. HeLa cells were transfected with the active GFP-RAB5 Q79L mutant and incubated with the inactive control CK-689 (A-C) or the selective ARP2/3 inhibitor, CK-666 (D-F). Cells were fixed and stained with phalloidin to detect filamentous actin (red). As depicted, there is a robust actin network at RAB5 Q79L endosomes in cells treated with CK-689 which is absent in cells treated with CK-666, suggesting that ARP2/3 is required for the actin nucleation.

**Expanded view 5.**
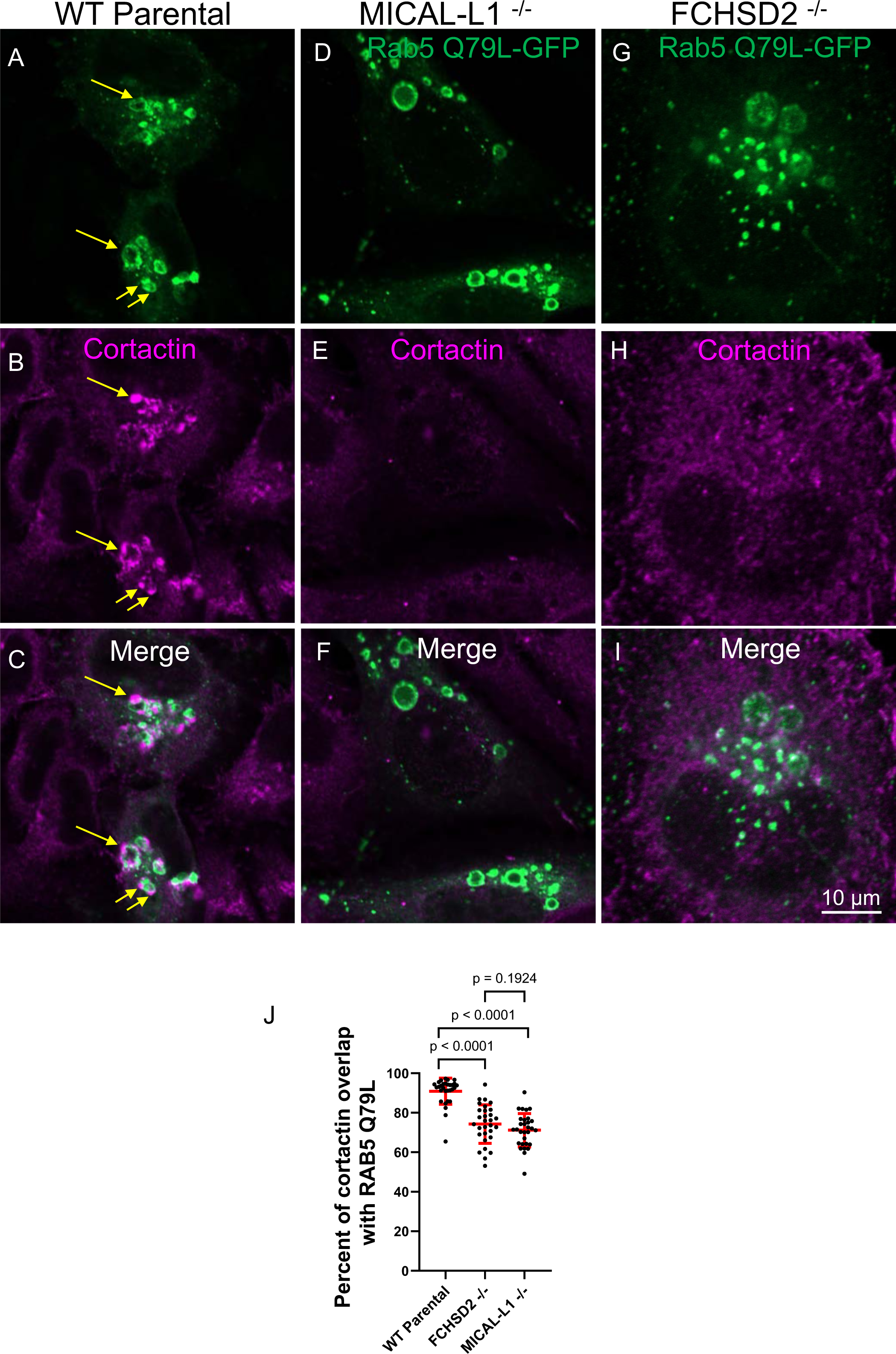
Both MICAL-L1 and FCHSD2 knock-out decrease branched actin localization at RAB5 Q79L endosomes. A-I. WT parental (A-C), MICAL-L1 knock-out (D-F), and FCHSD2 knock-out (G-I) HeLa cells were transfected with the active GFP-RAB5 Q79L mutant. Cells were fixed and immunostained with a primary antibody against cortactin to mark branched actin. WT parental cells show a robust branched actin network at the active RAB5 endosomes (A-C, yellow arrows), whereas both the MICAL-L1 knock-out (D-F) and FCHSD2 knock-out (G-I) show RAB5 endosomes that contain significantly less branched actin. J. Quantification of cortactin localization at RAB5 Q79L structures from A-I. Z-stack confocal images were captured and analyzed in Imaris by rendering 3D GFP-RAB5 Q79L structures. A region of interest around the RAB5 Q79L endosomes was demarked and the cortactin puncta within this region were also 3D rendered. Cortactin surfaces that contacted RAB5 structures were filtered by setting the maximal “shortest distance to surface” at 1x10^-7^ nm. The volume of cortactin structures contacting RAB5 structures was represented as a percentage of the total cortactin volume in the designated region.

